# Pitfalls of genotyping microbial communities with rapidly growing genome collections

**DOI:** 10.1101/2022.06.30.498336

**Authors:** Chunyu Zhao, Zhou Jason Shi, Katherine S. Pollard

## Abstract

Detecting genetic variants in metagenomic data is a priority for understanding the evolution, ecology, and functional characteristics of microbial communities. Many recent tools that perform this metagenotyping rely on aligning reads of unknown origin to a reference database of sequences from many species before calling variants. Using simulations designed to represent a wide range of scenarios, we demonstrate that diverse and closely related species both reduce the power and accuracy of reference-based metagenotyping. We identify multi-mapping reads as a prevalent source of errors and illustrate a tradeoff between retaining correct alignments versus limiting incorrect alignments, many of which map reads to the wrong species. Then we quantitatively evaluate several actionable mitigation strategies and review emerging methods with promise to further improve metagenotyping. These findings document a critical challenge that has come to light through the rapid growth of genome collections that push the limits of current alignment algorithms. Our results have implications beyond metagenotyping to the many tools in microbial genomics that depend upon accurate read mapping.

**HIGHLIGHTS:**

- Most microbial species are genetically diverse. Their single nucleotide variants can be genotyped using metagenomic data aligned to databases constructed from genome collections (“metagenotyping”).
- Microbial genome collections have grown and now contain many pairs of closely related species.
- Closely related species produce high-scoring but incorrect alignments while also reducing the uniqueness of correct alignments. Both cause metagenotype errors.
- This dilemma can be mitigated by leveraging paired-end reads, customizing databases to species detected in the sample, and adjusting post-alignment filters.

## Box 1: Glossary

**Conspecific.** Of the same species.

**Single nucleotide variant (SNV).** Nucleotide that differs between orthologous sites in conspecific genomes.

**Allele.** One of the observed sequences at a genomic locus, e.g., one of the nucleotides A, T, C, G at an SNV. Allele frequency is the proportional representation of one allele compared to others within a population.

**Genotyping.** Detecting genetic variants, identifying present allele(s), and quantifying allele frequencies, commonly performed through DNA sequencing.

**Metagenomics.** Shotgun sequencing of DNA extracted from a microbial community.

**Metagenotyping.** Genotyping with metagenomic data.

**Closely related genomes.** Genomes with genome-wide similarity above a threshold (e.g., 92% average nucleotide identity).

**Closely related species.** Species that share at least one pair of closely related genomes, may be estimated with one representative genome per species for computational efficiency.

**Reference bias.** Reduced alignment accuracy between divergent genomes of the same species. In the context of metagenotyping, genetic differences between genomes in the sample versus database affect the rate and accuracy with which reads can be aligned.

**Uniquely-mapping.** Sequencing read with one reported alignment or a best alignment that scores much higher than the second-best alignment.

**Multi-mapping.** Sequencing read with two or more different alignments reported.

**Cross-mapping.** Sequencing read from one species aligned to another species. Also known as an off-target alignment.

**On-target alignment.** Sequencing read aligned to the correct species.

**Post-alignment filter.** Rule used to discard alignments, for example, based on sequence similarity or uniqueness.

**MAPID.** Sequence identity between a read and the database sequence to which it is aligned.

**MAPQ.** A measure of alignment uniqueness based on the ratio of alignment score of the best versus second best alignment.

**Vertical coverage.** Number of reads aligned at a nucleotide or other genomic element.

**Horizontal coverage.** Proportion of nucleotides in a genome that are covered by alignments (e.g., at least two aligned reads).

**Paired-end.** Sequencing strategy in which both ends of a molecule are sequenced. In the context of metagenomics, both reads in a pair should be aligned nearby and to the same species.

**K-mer.** A nucleotide sequence of length k, where k is typically a small integer.

## INTRODUCTION

Most microbes harbor immense within-species genetic variation, with strains differing in terms of single nucleotides, gene copy number, and genome organization. Recognizing genetic differences between conspecific genomes is important for many reasons. First, genetic variation can have functional consequences, ranging from differential metabolism to acquisition of pathogenicity and antibiotic resistance (Chattopadhyay et al., 2009; Leshem et al., 2020; Maini Rekdal et al., 2019; Zeng et al., 2019). Genotype atlases enable association studies that promise to reveal many such genotype-phenotype links (Power et al., 2017). Second, variants are useful markers for tracking strains and mobile genetic elements, allowing investigations into their ecological dynamics (Saak et al., 2020; Smillie et al., 2018). For human-associated microbes, this enables epidemiological studies of clinically important strains, including transmission and dispersal of different lineages (Mitchell et al., 2020). Finally, genetic variants may be utilized to infer the evolutionary forces acting on microbial species, shedding light on the roles of drift, selection, and recombination across taxonomic groups and environments (Garud and Pollard, 2020; Shoemaker et al., 2022; Van Rossum et al., 2020). Thus, there is great interest in characterizing the genetic diversity of microbiomes beyond the species level.

While cultured isolates have been genotyped for decades, metagenotyping–genotyping species using shotgun metagenomic DNA sequences–is greatly expanding the field of microbial population genetics by enabling researchers to detect genetic variation at an unprecedented scale and in new settings (Garud and Pollard, 2020; Shoemaker et al., 2022; Van Rossum et al., 2020). Key benefits include being able to capture genetic variation across whole genomes, in uncultured species, in samples that are ecologically diverse, and for many species in parallel with a single experiment. Metagenotyping many species from a complex community sampled from its natural environment not only enables direct investigation into microbe-environment associations at a precise taxonomic resolution, but it also reveals co-occurring and co-excluding lineages. This may include interactions between strains of the same species as well as inter-species relationships, such as strain-specific phage resistance or bacteria-fungi associations (Forbes et al., 2018).

Many bioinformatics pipelines have been developed for metagenotyping (reviewed in (Ghazi et al., 2022)). Most of these methods are reference-based (**Table S1**), meaning they use alignment algorithms to map metagenomic reads to a database of genomes or gene sequences and apply established genotyping workflows to call variants for each species. Commonly used aligners include Bowtie2 (Langmead and Salzberg, 2012), BWA (Li and Durbin, 2009), and minimap2 (Li, 2018). Metagenotyping suffers from many problems previously documented in the context of genotyping individual strains, ranging from sequencing errors and alignment errors to reference bias (Anyansi et al., 2020; Bush et al., 2020; Ghazi et al., 2022). However, these known errors are amplified by the massive diversity and high proportion of low-abundance species present in metagenomic data, coupled with using multi-species reference databases. These factors also present several unique challenges, including cross-mapping of reads to the wrong species (Hovhannisyan et al., 2020) and reduced alignment uniqueness (Zhao et al., 2022). The core issue is that metagenotyping tools utilize alignment algorithms in scenarios that are more complex than the contexts for which the aligners were developed (**Table 1**).

**Table 1.** Complexity of sample and database across genotyping contexts.

|  | Example | Species in Sample | Intra-species Variation per Species in Sample | Species in Database |
| --- | --- | --- | --- | --- |
| **Isolate genotyping** | Clonal isolate culture | One | None - Low | One |
| **Community genotyping** | In vitro evolution experiment | One or Few | Low - Medium | Same as Sample |
| **Metagenotyping** | Shotgun sequencing of a stool sample | Many | Low - High | Many |

In this Synthesis, we highlight the challenges that arise in reference-based metagenotyping of bacterial single-nucleotide variants (SNVs) in short reads, which is currently the most common strategy for studying genetic variation in microbial communities. We focus on problems that are exacerbated by or unique to metagenotyping as compared to genotyping a single isolate. Through surveying current genome databases, we show that an increasing number of bacterial lineages contain sequences for multiple closely related species as well as a growing diversity of genomes per species. These changes have benefits, but also some unfortunate downsides. Through examples and simulations, we explore why metagenotyping errors tend to be worse in lineages with many closely related species and/or high intraspecific diversity. Next, we evaluate post-alignment filters and custom genome databases as potential solutions to combat metagenotyping errors. We then review ongoing and future work that could further improve the accuracy and utility of SNV metagenotypes, including alternatives to current alignment algorithms. Recognizing that this field is evolving quickly, we discuss metagenotyping for other variant types, taxonomic groups, and sequencing technologies. We conclude with a review of alternatives to standard alignment algorithms and broader implications of our findings for the use of alignment in microbiome research.

## SOURCES OF GENETIC VARIATION IN METAGENOMES

Each metagenomic sample contains reads from many species. Sequencing reads from orthologous regions of any one of these species may harbor nucleotide differences due to genetic diversity captured in the sample (**Figure 1A**) as well as sequencing errors. Genetic diversity has multiple sources (Ghazi et al., 2022; Shoemaker et al., 2022; Van Rossum et al., 2020). One common source is the presence of two or more divergent lineages of the same species within a metagenome. Any lineage abundant enough to be sequenced will contribute to the presence and allele frequencies of within-sample SNVs. When the lineages are not closely related, many SNVs will be detected genome-wide. Another way to generate genetic variation is a new mutation arising within a clonal lineage. If the mutation becomes prevalent enough in the community to be captured and sequenced, it will be detected as a within-sample SNV. Horizontal gene transfer and homologous recombination also introduce SNVs. Metagenotyping aims to detect all these sources of within-species genetic variation.

**Figure 1.**
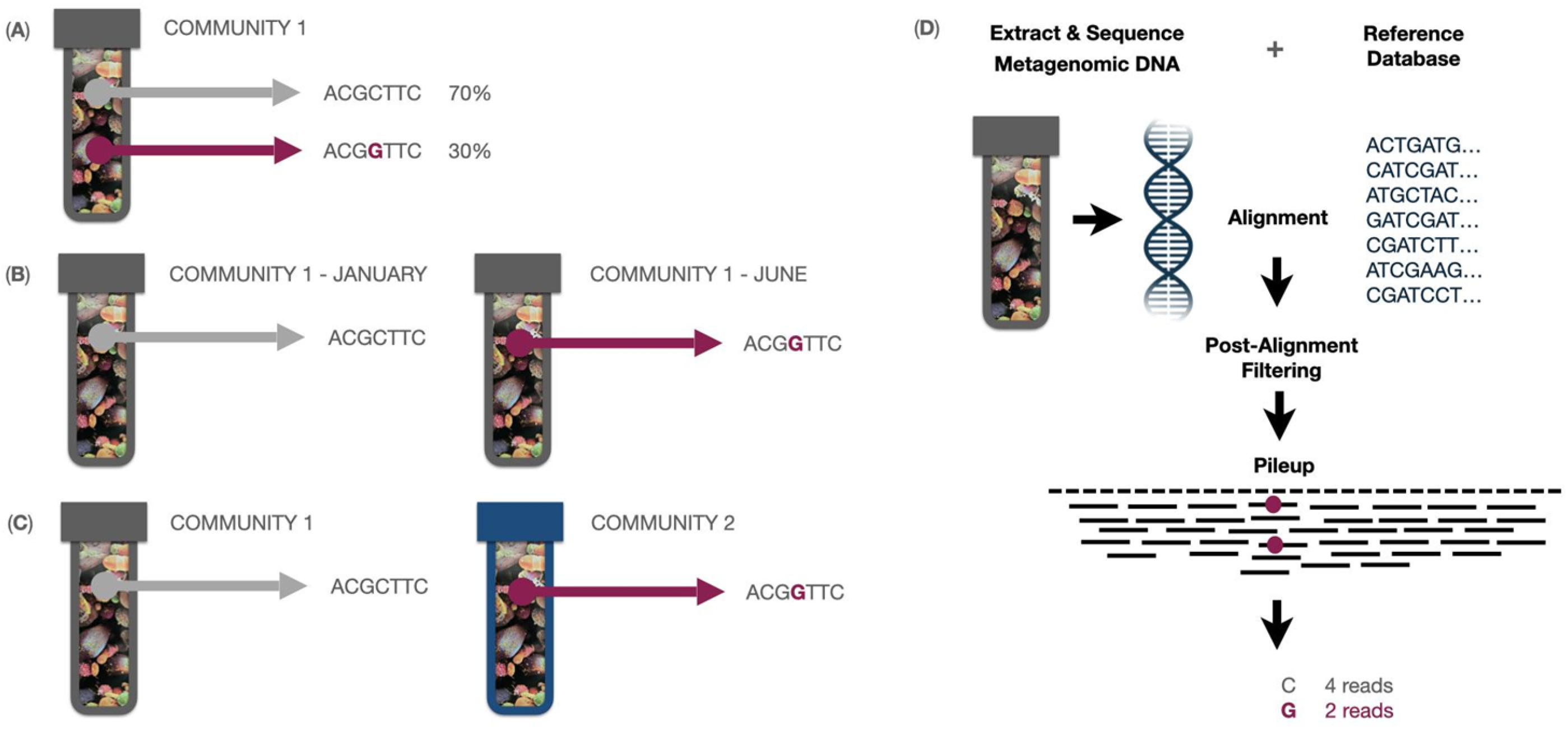
Detecting SNVs in microbiome samples. Within-species genetic variation, such as single nucleotide variants (SNVs), can be detected within and between communities using metagenotyping. (**A**) A single community may harbor multiple strains of the same species at the same time. A biallelic SNV is shown. The C allele is present in 70% of genomes, while the other 30% have the G allele. This genetic variant may be detected by metagenotyping a single sample. (**B-C**) Across-sample analysis of metagenotypes may reveal population SNVs. (**B**) Over time, conspecific genomes in a community develop genetic differences through strain replacement, recombination, horizontal gene transfer, and de novo mutation. (**C**) Two communities may harbor divergent strains with many SNVs. (**D**) Reference-based metagenotyping involves mapping metagenoic reads to a reference database of representative genome sequences (or marker genes) using alignment or an alternative method, such as k-mer exact matching. Typically, alignments are filtered before detecting SNVs in the pileup of aligned reads. The detected SNV alleles and their read counts are output to a file. Across-sample analysis involves merging this data for a set of samples.

Metagenomic sequencing reads from an orthologous region of the same species typically harbor multiple nucleotide differences when we compare samples from the same community over time (**Figure 1B**) or from different communities (**Figure 1C**). Conserved nucleotides may represent recently acquired mobile elements, sites with strong negative selection, or closely related lineages (when genome-wide). Some metagenotyping tools merge alleles of SNVs detected within a set of samples to enable across-sample genetic analyses and to identify population SNVs detected in more than one sample (Olm et al., 2021; Schloissnig et al., 2013; Van Rossum et al., 2021; Zhao et al., 2022). In other cases, users need to write customized scripts for cross-sample metagenotype analysis (Shi et al., 2022). The SNV merging step may only use the consensus allele for each sample (i.e., the nucleotide observed in the most reads) or it can preserve within-sample variation by including the read counts for each nucleotide or for the two most frequent ones. With the resulting population SNVs for each species in a set of metagenomic samples, researchers can perform a rich array of analyses, including strain deconvolution or haplotype inference, metagenome-wide association studies, and tests for positive selection across gene families, as reviewed in-depth elsewhere (Ghazi et al., 2022). All of these downstream investigations depend upon accurate metagenotypes.

## STEPS INVOLVED IN REFERENCE-BASED METAGENOTYPING

To understand when metagenotyping breaks down, it is important to know how metagenotypes are generated (**Figure 1D**). All metagenotyping starts with a DNA sequencing library generated from a microbial community. Reads may be quality controlled, trimmed, or taxonomically filtered (e.g., to remove host reads or contaminants). Next, reference-based methods (**Table S1**) employ alignment algorithms to map each read to the putative species and genome coordinates from which it was derived. This is done using a multi-species database, typically consisting of one representative genome for each distinct species. Each alignment is scored based on similarity of the read to the database sequence after taking into account base quality, and these scores are used to assess the uniqueness of the best alignment (e.g., MAPQ in Bowtie2, which compares the scores from the best and second-best alignment). In a post-processing step, potentially erroneous alignments are filtered out based on the alignment score and uniqueness. Then, the pileup of remaining reads is used to detect SNVs, producing a metagenotype vector for each species in each sample. Some software packages additionally quantify allele frequencies, while others use pileups to metagenotype structural variants of various sizes (Greenblum et al., 2015; Zeevi et al., 2019). Although metagenotyping appears to be a straightforward extension of well-vetted genotyping methodologies, the complexity of the sequencing library and the multi-species database create some unique challenges for alignment algorithms and post-alignment filtering.

## ALIGNMENT PITFALLS & THEIR EFFECTS ON METAGENOTYPING

Accurate alignment is critical for generating a correct metagenotype. In this study, we say the species from which a read was derived is the on-target species, and all others are off-target species. Let us consider all the possible outcomes when aligning a sequencing read to the on-target genome (**Figure 2A**), ignoring at first the huge diversity of species in a metagenotyping database. In the best-case scenario, the read has one high-scoring alignment, so it is retained for pileup. We have high confidence in nucleotide differences from the reference genome, because we trust that the read is correctly aligned. As uniqueness decreases, the probability that the best alignment is correct decreases (JohnUrbanGenome, 2022). If uniqueness gets too low, we say that the read is multi-mapping, as in other genomics applications (Deschamps-Francoeur et al., 2020; Zheng et al., 2019), and it will not be retained for the pileup. The read will not be aligned at all when the best alignment’s score is low. These filters help to prevent false positive SNV calls. On the other hand, an alignment score threshold will also remove correctly aligned reads from strains that are diverged from the reference genome. This phenomenon is known as reference bias (Garrison et al., 2018), and it contributes to false negative SNVs. Uniqueness filtering also results in false negative SNVs, and it can bias allele frequency estimates. Thus, metagenotyping methods face a tension between controlling erroneous SNV calls and ensuring that true SNVs are detected.

**Figure 2.**
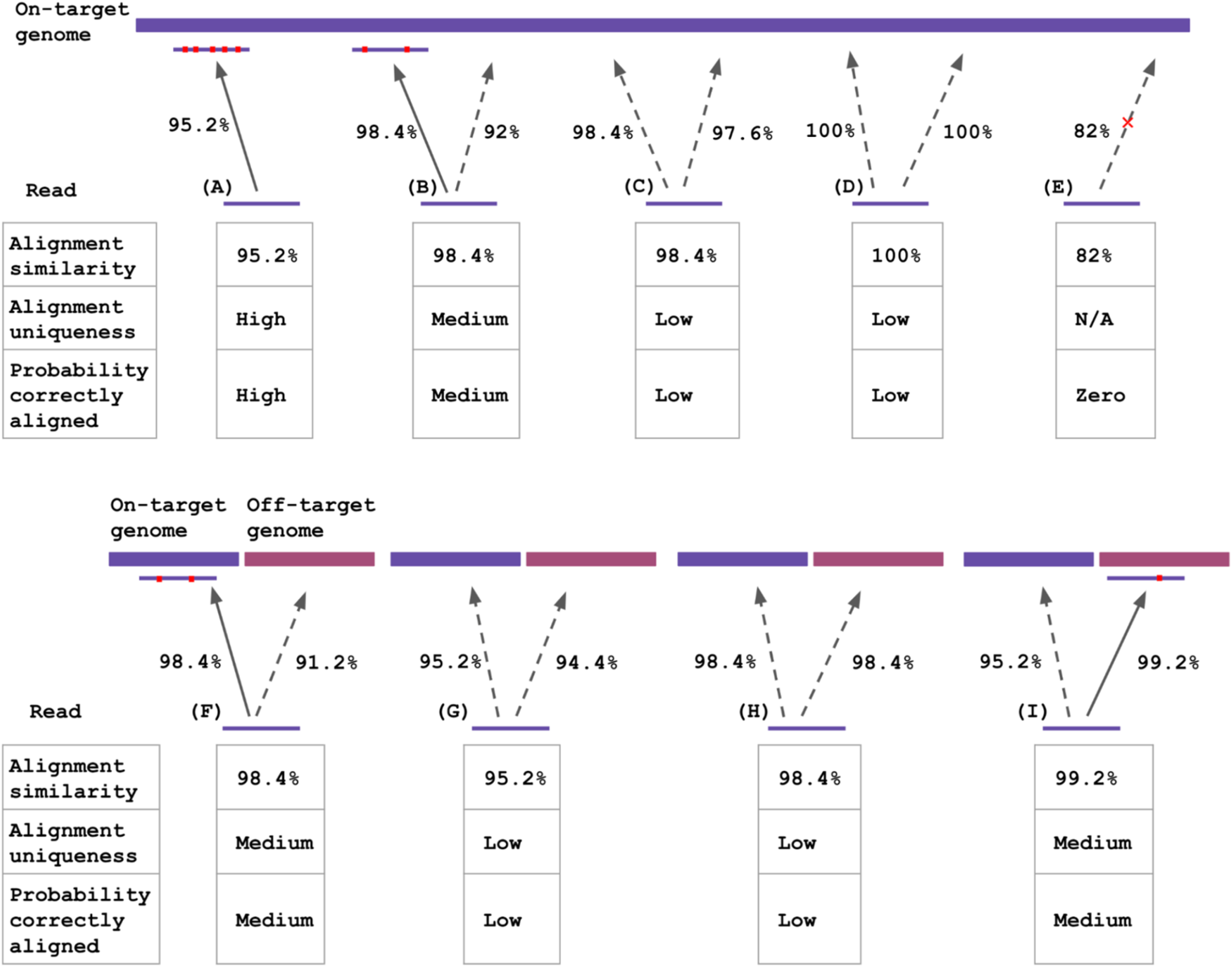
Common alignment pitfalls in the context of reference-based metagenotyping. Possible outcomes of aligning one read from a metagenomic sequencing library to a metagenotyping database. Alignment similarity, alignment uniqueness, and the probability that the read is correctly aligned after applying post-alignment filtering is recorded below each example. Post-alignment filtering: The best alignment is retained if sequence similarity between the read and the genome is high enough and if the alignment is unique enough compared to the second best alignment. Solid arrow: the best alignment, passes post-alignment filtering; Dashed arrow: read aligns but is not the best alignment and/or fails post-alignment filtering. Arrow with cross: read does not align, best alignment is below the aligner’s minimum score. Alignment similarity indicated on all arrows. For aligned reads, SNVs between the read and the genome are indicated by red dots. **Top:** The database contains only an on-target genome, which is a different strain of the same species as the read (purple). (**A**) Uniquely aligned read: aligned one place with high similarity and uniqueness, so it passes post-alignment filtering. It is likely a correct alignment. (**B**) Aligned read with low uniqueness: the second best alignment is pretty good, so it may fail post-alignment filtering depending on the uniqueness threshold. Confidence in the best alignment is reduced. (**C**) Multi-mapping read: aligned two places and the best alignment has only slightly higher similarity, so it will probably fail post-alignment filtering. (**D**) Perfectly multi-mapping read: aligned two places with identical similarity, so it will fail post-alignment filtering. (**E**) Unaligned read: no alignments reported. **Bottom:** The database contains the on-target genome plus an off-target genome of another species (maroon). (**F**) Aligned read with low uniqueness: the second best alignment is to the off-target species but has fairly high similarity. The uniqueness threshold will determine if it passes post-alignment filtering. (**G-H**) Multi-species multi-mapping reads. (**I**) Cross-mapping read: the best alignment is off-target. It may pass post-alignment filtering, depending on the uniqueness threshold. A higher uniqueness threshold would reduce cross-mapping and false positive SNVs in the off-target species, but this would also eliminate correct alignments as in **B** and **F**. Filtered out and unaligned reads can bias SNV metagenotypes in the on-target species.

Read alignment is even more challenging in reality, because both the metagenomic sample and the database contain many species (**Figure 2B**). It is not known a priori which reads in the sequencing library come from which species. Furthermore, short reads may have high-scoring alignments to genomes from off-target species, a problem known as cross-mapping (Hovhannisyan et al., 2020). It can occur between highly conserved and horizontally transferred sequences in distantly related species, but it affects many genomic loci when the database contains closely related species. The key issue is that homologous sequences from the representative genomes of on-target and off-target species in the database compete for reads. If neither alignment is unique enough, the read is filtered out. This affects reads that carry nucleotide differences from the on-target representative genome and those that do not, contributing to errors in SNV detection and allele frequency estimates. Another source of error is when the off-target genome has the best alignment with sufficiently high uniqueness for the read to be retained. This affects the metagenotypes of both species, and it is more frequent with reads carrying nucleotide differences from the on-target genome.

Clearly, reference bias, multi-mapping, and cross-mapping have the potential to negatively impact metagenotype accuracy. We next look at how widespread these problems are across bacteria. Then we quantify their effects on metagenotypes and use these results to explore two ways to combat the problem: changing post-alignment filtering thresholds and customizing genomes in the database to be as similar as possible to the metagenomic sample.

## RAPID GROWTH IN BACTERIAL GENOME DATABASES

Starting around 2015, the number of species with at least one genome sequence began to skyrocket (**Figure 3A**), with hundreds of thousands of prokaryotic species now represented in NCBI Assembly (Kitts et al., 2016), European Nucleotide Archive (Leinonen et al., 2011), and other databases (Chen et al., 2021a). This explosion of genomes is driven in part by lower costs and higher throughput of DNA sequencing, coupled with algorithms for assembling isolate genomes from short reads. Meanwhile, culture collections have grown rapidly due to technological advances and concerted efforts to capture difficult to grow strains from diverse environments (Groussin et al., 2021; Mukherjee et al., 2017; Nowrotek et al., 2019; Sarhan et al., 2019; Sood et al., 2021). Another major source of genomes has been assemblies generated directly from tens of thousands of metagenomic sequencing libraries sampled from diverse environments (Levin et al., 2021; Parks et al., 2017). These metagenome assembled genomes (MAGs) comprise a large proportion of databases such as GEM (natural environments) (Nayfach et al., 2021) and UHGG (human gut) (Almeida et al., 2021b) (**Figure 3B**). MAGs have been particularly useful for capturing genomes from species that are difficult to isolate with traditional culturing techniques. Species that are medically important, laboratory models, and prevalent in environments that receive the most research attention are over represented in genome databases, though these biases are decreasing somewhat.

**Figure 3.**
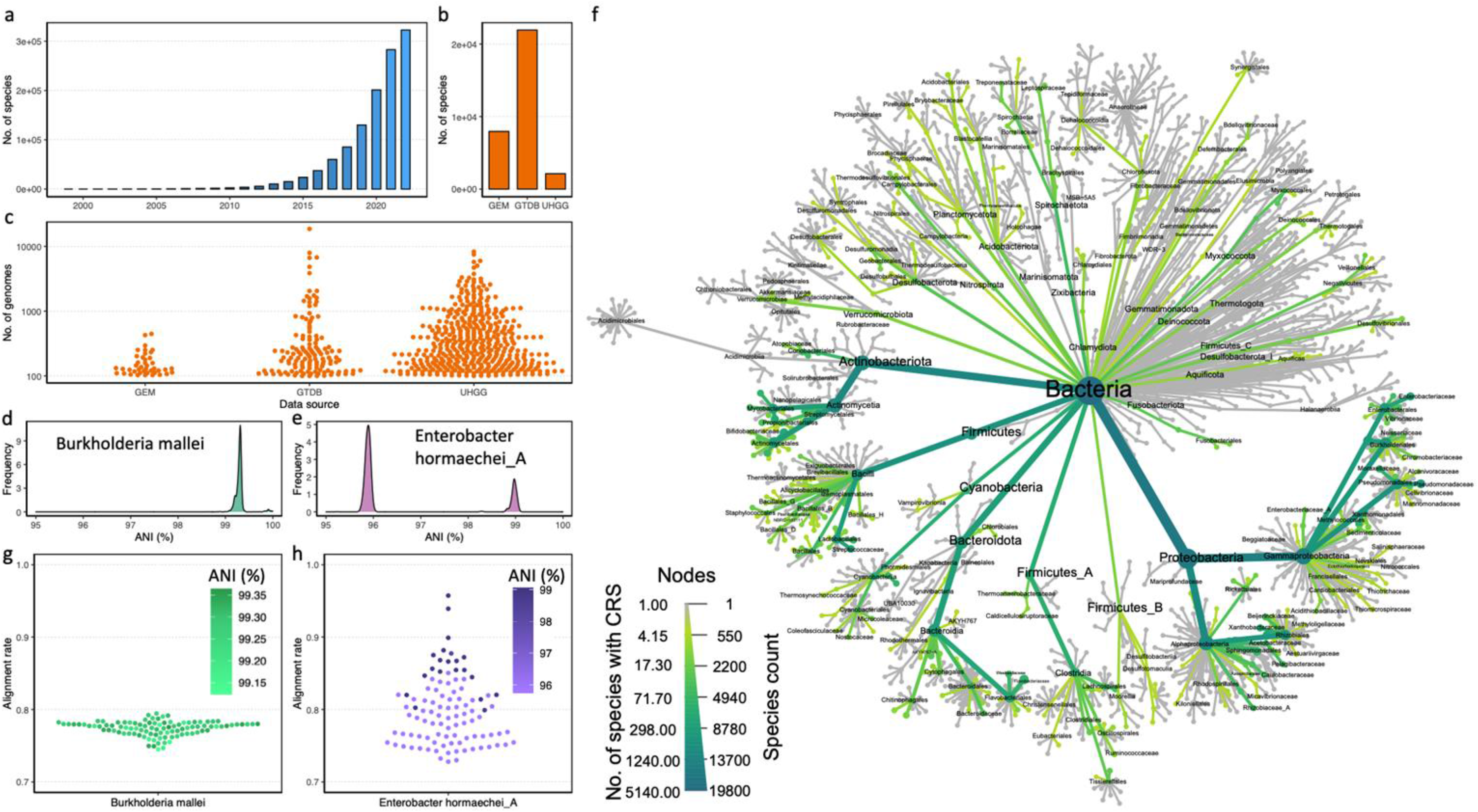
Rapid growth in prokaryotic genome sequences uncovers diverse and closely related species. (**A**) The number of prokaryotic species with at least one genome assembly has grown rapidly in recent years. NCBI Assembly database (as of May 2022) by year from 1999 to 2022. (**B**) Many large genome collections are available today. Shown are Genomes of Earth’s Microbiomes (GEM; June 1, 2022), Genome Taxonomy Database (GTDB; version R207), and Unified Human Gastrointestinal Genome collection (UHGG; v1.0). (**C**) It is now common for a prokaryotic species to have multiple genome sequences. (**D-E**) Species differ in the amount of intra-specific genetic diversity in genome databases. Genome-wide average nucleotide identity (ANI) between the representative genome and conspecific genomes is shown for two example species chosen from GTDB to represent different population structures at the extremes of intra-species ANI. (**D**) *Burkholderia mallei* (RS_GCF_000011705.1). (**E**) *Enterobacter hormaechei_A* (RS_GCF_001729745.1). (**F**) A heat-tree (cladogram) showing the prevalence of bacterial taxa with a closely related species (CRS), defined as 92%-95% identity (1 - Mash distance, a fast approximation of ANI). Species with CRS are most common in Proteobacteria and specific lineages of Actinobacteria. This phylogenetic distribution in part reflects the large amount of sequencing and assembly effort focused on pathogens, model organisms, and the human microbiome. (**G**) We simulated metagenomic reads from 100 *B. mallei* genomes at 30X coverage and aligned them to the GTDB *B. mallei* representative genome. Alignment rate is highest when reads come from a genome that has higher ANI with the representative genome. (**H**) *E. hormaechei_A* shows the same relationship between alignment rate and ANI but with more variability in both variables.

One major benefit of more species with genomes is that a greater diversity of species can be metagenotyped with reference-based methods. For example, UHGG provides reference genomes for nearly all prevalent prokaryotic species residing in the stool of individuals from North America or Europe (Almeida et al., 2021b). Using samples from the PREDICT cohort (Asnicar et al., 2021), we estimate that this translates to an alignment rate of ~80%, which is an improvement over older databases (e.g., ~65% alignment rate for NCBI in 2013). Database coverage is unfortunately lower but also improving for other human populations (Smits et al., 2017) and other environments (Nayfach et al., 2021). It is important to keep in mind that new genomes are only helpful to metagenotyping tools if their quality is high enough to generate accurate genotypes. When reference genomes are fragmented, incomplete or contaminated with sequences from off-target species, the number of reads that can be correctly mapped is reduced and fewer sites can be metagenotyped (Olm et al., 2021; Shi et al., 2022; Van Rossum et al., 2021; Zhao et al., 2022).

In parallel with increasing the species diversity of genome databases, recent sequencing and assembly efforts have also greatly expanded the number of genomes per species. Genomes are typically grouped into species using algorithms based on sequence similarity. Genome-wide average nucleotide identity (ANI) greater than 95% serves as an operational species definition, though this threshold is debated and not strictly followed (Jain et al., 2018; Murray et al., 2021; Olm et al., 2020; Rodriguez et al., 2021). There are now dozens of species with more than 1,000 genome sequences (**Figure 3C**). For many species, the genomes are highly clonal. For example, *Burkholderia mallei* has 1,698 genomes in GTDB, and most pairs of genomes are ~99% identical (**Figure 3D**). Other species show a greater range of sequence diversity, and some species display population genetic structure with multiple clusters of genomes. Pairs of *Enterobacter hormaechei_A* genomes in GTDB, for instance, tend to have either ~99% ANI or ~96% ANI, reflecting two divergent lineages (**Figure 3E**).

With more species in genome databases and more genomes per species, boundaries between species are getting closer together and in some cases blurred (Jain et al., 2018; Murray et al., 2021; Olm et al., 2020; Rodriguez et al., 2021). Thousands of bacteria now have a closely related species with >92% ANI (**Figure 3F**). Closely related species occur in specific lineages of most phyla, and they are most numerous in Proteobacteria and Actinobacteria. Coupling closely related species with divergent lineages of the same species (close to 95% ANI), it is possible that a strain of one species is more similar to a genome from a closely related species than one from its own species, or at least that some parts of its genome are. In some lineages, new genomes are basically filling the gaps in sequence similarity between species that were previously separated. Clearly, the growth in genome sequencing has consequences for accurate read alignment to reference genomes in metagenotyping tools.

## MEASURING THE EFFECTS OF DATABASE GROWTH ON METAGENOTYPING

To quantify how closely related genome sequences affect metagenotypes, we performed a series of carefully controlled *in silico* experiments in which metagenomic sequencing reads were generated from genomes using the read simulator ART (Huang et al., 2012) and aligned with Bowtie2 (Langmead et al., 2019) to databases in which we vary the following parameters using fastANI (Jain et al., 2018): (1) inter-species ANI of the closest off-target genome and (2) intra-species ANI between the genome from which reads were simulated and the on-target representative genome. Since these were simulations, we could directly track alignment rates (% aligned, horizontal coverage, vertical coverage), cross-mapping rates (% aligned reads that are aligned to an off-target genome), and SNV accuracy (precision, recall) for reference versus alternative alleles. The reference allele is the nucleotide matching the database sequence, and all other nucleotides are alternative alleles. We first evaluated the full spectrum of errors with no post-alignment filters (alignment parameters: *bowtie2 --no-unal -X 1000.0 --end-to-end --very-sensitive*), and then we examined the effects of applying various filters. We varied simulated read coverage and observed a plateau in performance statistics starting around 10X (**Figure S1**). Trends in all measurements were qualitatively similar across coverage levels. We therefore used 20X in several of our analyses to demonstrate a specific problem or solutions. We emphasize that the identified problems are expected to be even worse in complex microbial communities where many species are at lower coverage values, and we refer readers to **Figure S1** for results across coverage levels. By repeating this workflow for hundreds of bacterial species with different population structure, diversity, and distance to closely related species, we captured a huge variety of scenarios. These analyses vividly illustrated the hypothetical alignment problems from **Figure 2** using millions of reads, and they enabled us to evaluate potential solutions in a quantitative manner.

## INTRA-SPECIES DIVERSITY BIASES METAGENOTYPES TOWARDS REFERENCE ALLELES

Reference bias is well known to affect genome comparisons (Bush et al., 2020; Garrison et al., 2018; Gunther and Nettelblad, 2019). Not surprisingly, our simulations showed that it is also at play in metagenotyping workflows. Using alignment of metagenomic reads simulated from 100 conspecific genomes of *B. mallei* (**Figure 3G**) or *E. hormaechei_A* (**Figure 3H**) to a single representative genome of each species as examples, we observe a clear correlation between alignment rate and genome-wide ANI to the representative genome. Repeating this analysis with all high-quality UHGG genomes for 327 species that have intra-species ANI ranging between 95 and 100%, we observed a significant positive correlation between alignment rate and ANI (**Figure 4A**). As intra-species ANI approaches the species boundary, only ~75% of reads are aligned on average. Importantly for metagenotyping, the probability of a correct alignment is lower for reads with differences from the reference genome. Hence, both precision (**Figure 4B**) and recall (**Figure 4C**) are lower for SNV metagenotypes of alternative versus reference alleles. Thus, when the representative genome in a metagenotyping database is diverged from the genome(s) in a metagenomic study, fewer SNVs can be metagenotyped and the accuracy of the allele frequencies at SNVs that are metagenotyped will be biased towards the reference allele.

**Figure 4.**
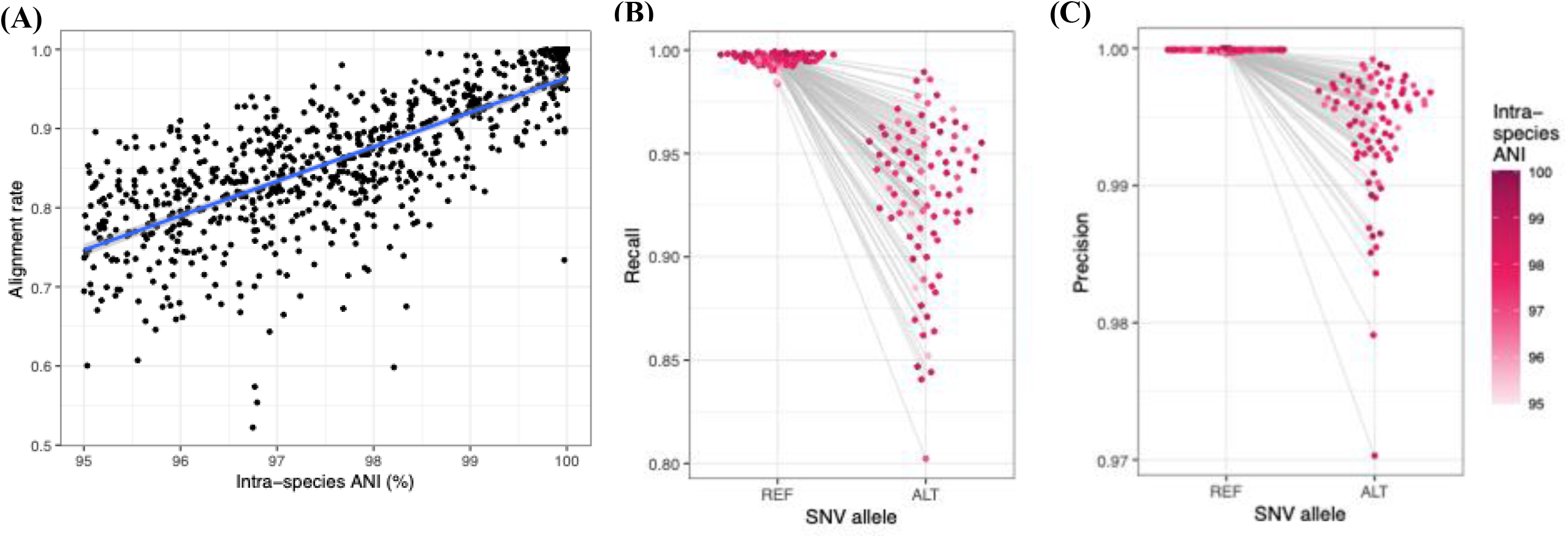
Reference bias reduces the number and accuracy of metagenotyped SNVs for divergent lineages. (**A**) We quantified reference bias for 327 species selected from the UHGG database to represent a range of different levels of intra-species diversity. Simulated metagenomic reads from all genomes of each species were aligned to a single high-quality representative genome, one species at a time with no other genomes in the Bowtie2 database. Across species, alignment rate is positively correlated with ANI (Pearson’s correlation 0.75, p<2.0×10^-16^) and ranges from 52% to 100%. Alignment rate is computed as the fraction of aligned reads over total simulated read counts. (**B-C**) We metagenotyped SNVs in alignments from (**A**) for reads simulated from 86 high-quality NCBI genomes, without any post-alignment filtering. SNV recall and precision are correlated with intra-species ANI (i.e., similarity of the NCBI genome in the metagenomes to the UHGG genome in the database) for reference (REF) alleles. (**B**) As expected, unaligned reads tend to carry more alternative (ALT) alleles, and hence recall is notably higher for REF alleles. (**C**) Precision is very high for REF alleles and somewhat lower for ALT alleles.

## CROSS-MAPPING IS PREVALENT IN METAGENOTYPING WORKFLOWS

We next used our simulation framework to quantify cross-mapping of metagenomic reads to a genome from the wrong species. To do so, we repeated the analyses described above with the addition of a second, off-target genome in the database. For each species, we iterated through a set of off-target genomes ranging from closely related species (inter-species ANI ~95%) to more distantly related species (inter-species ANI <77%). In the absence of cross-mapping, no reads should align to the off-target genome.

This analysis showed that cross-mapping is prevalent and increases in frequency as the off-target genome approaches the species boundary (**Figure 5A**). It is also bi-directional, meaning that from the perspective of the on-target species, similar amounts of reads are lost to and stolen by closely related species (**Figure S2**). On average, sites of the off-target genome have ~10X vertical coverage in simulations with 20X coverage (**Figure S3**), indicating that erroneous alignments are not limited to a small number of reads from a genomic locus (e.g., those with a particular sequencing error). We initially thought that cross-mapped reads might therefore be piling up in specific extremely conserved loci. However, we found that closely related off-target species (ANI > 92%) have a median horizontal coverage of 23.5% (range 3.8%-71.3%), which drops down to ~5% for more distant off-target species (**Figure 5B**). This positive relationship between ANI and horizontal coverage is consistent with prior findings based on genomes (Olm et al., 2020). Together, our results show that cross-mapping occurs broadly across the genomes of closely related species and may affect a substantial proportion of reads.

**Figure 5.**
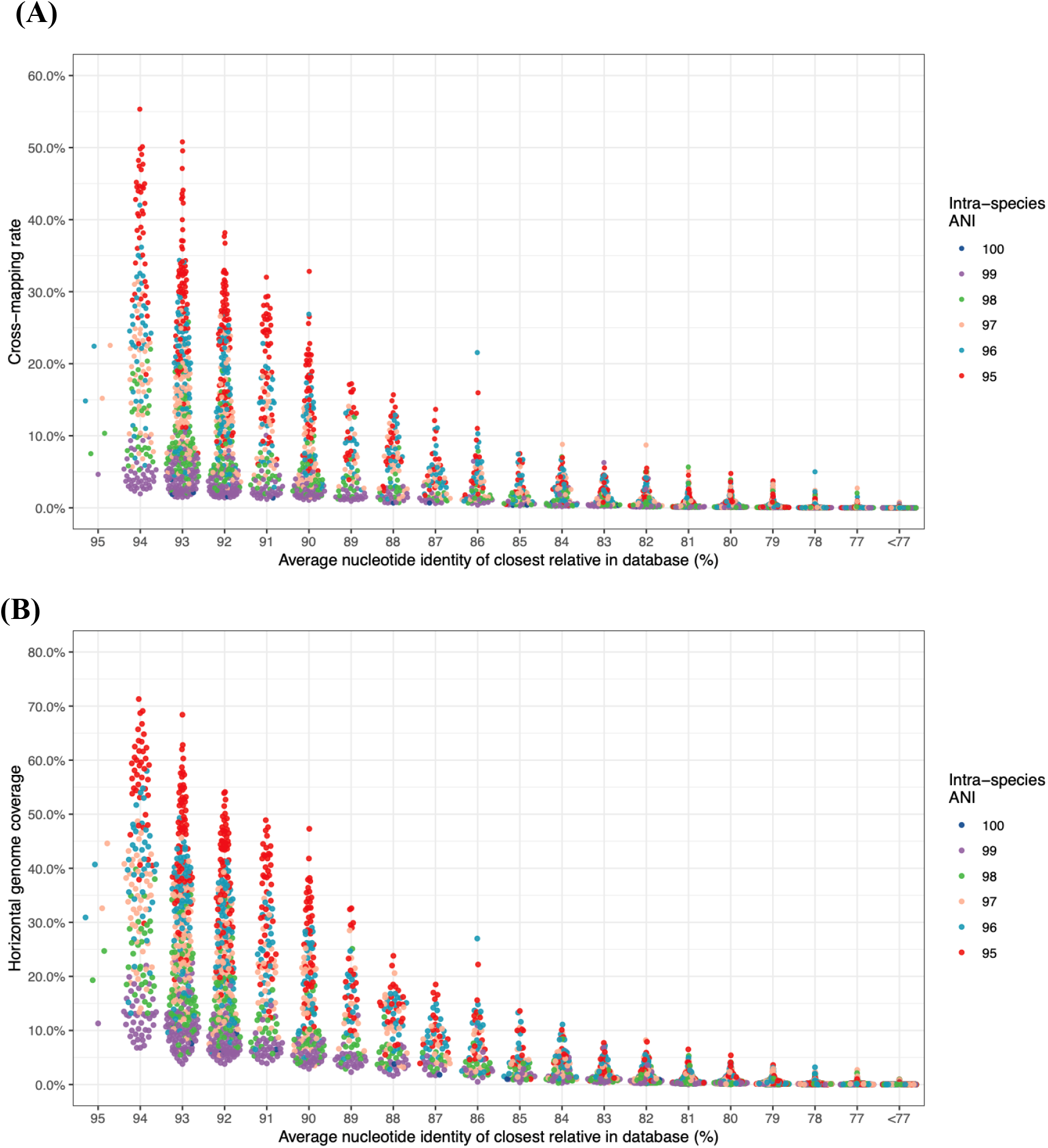
Cross-mapping is prevalent. Using UHGG, reads were simulated at 20X coverage from one genome and aligned to a database containing two representative genomes: the on-target species (color: ANI to the simulation template) and an off-target species (horizontal axis: inter-species ANI). All aligned reads were retained (no post-alignment filtering). (**A**) We observe increased cross-mapping as the template genome gets more diverged from the representative genome and as the two species become more closely related. Cross-mapping rate is the proportion of all aligned reads incorrectly mapped to the off-target genome. (**B**) Crossmapped reads can cover a high proportion of the off-target genome, with similar trends as the crossmapping rate. Horizontal coverage is the proportion of nucleotides in the off-target genome covered by at least two reads.

Within these trends, cross-mapping varies quite a bit by species and even within the same species. Both the cross-mapping rate (**Figure 5A**) and the horizontal coverage of the off-target genome (**Figure 5B**) tend to be higher when reads are simulated from a species with greater intra-species diversity, after controlling for inter-species ANI. In the worst-case scenario when intra-species ANI is ~95% and there is a closely related off-target genome in the database (~94% interspecies ANI), as much as ~50% of reads can be aligned to the wrong genome, and the off-target genome may have up to ~70% horizontal coverage. But when intra-species diversity is lower and the off-target genome is less closely related, we observe less cross-mapping (median 3% for all off-target genomes having inter-species ANI ≥ 92%) and lower off-target horizontal coverage (median 9.2%). Because these results were generated with actual pairs of genomes present in metagenotyping databases, we conclude that closely related species drive a great deal of cross-mapping, especially in lineages with diverse species where species boundaries are blurred and when the representative genome for the on-target species is diverged from the strains in the metagenome. Since there is relatively little cross-mapping with more distantly related species, we focus in the following simulations on one closely related off-target species for each on-target species.

## LOW ALIGNMENT UNIQUENESS IS A MAJOR DRIVER OF ERRONEOUS METAGENOTYPES

We next used our Bowtie2 alignment results for simulated metagenomic reads from *Collinsella sp003458415* against databases with different degrees of closely related species to examine the fate of reads without any post-alignment filters. We began by looking at the distribution of MAPID and MAPQ values across scenarios. MAPID is the alignment sequence identity, and it is used to evaluate the match between metagenomic read and reference genome. MAPQ measures how much better the highest-scoring alignment is compared to the second-best alignment. A high MAPQ reflects a unique alignment, and this statistic is used to decide how confident one is in the reported alignment.

When only the on-target *C. sp003458415* genome is in the database, most alignments have high MAPQ (mean: ~40, 95th percentile: ~25; **Figure 6A**) and MAPID > 95% (**Figure 6B**). However, we observed high variability in both MAPID and MAPQ. The low MAPID values reflect alignments where the genome from which reads were simulated differs from the reference genome (reference bias). This is expected based on our prior results, because *C. sp003458415* is a diverse species (intra-species ANI = 95.16%). The reads with low MAPQ show that multi-mapping occurs within this species. Thus, with a single genome of the correct species in the database, MAPID and MAPQ are performing as expected. They detect alignments where read mapping is uncertain due to divergence from the reference genome or paralogous sequences.

**Figure 6.**
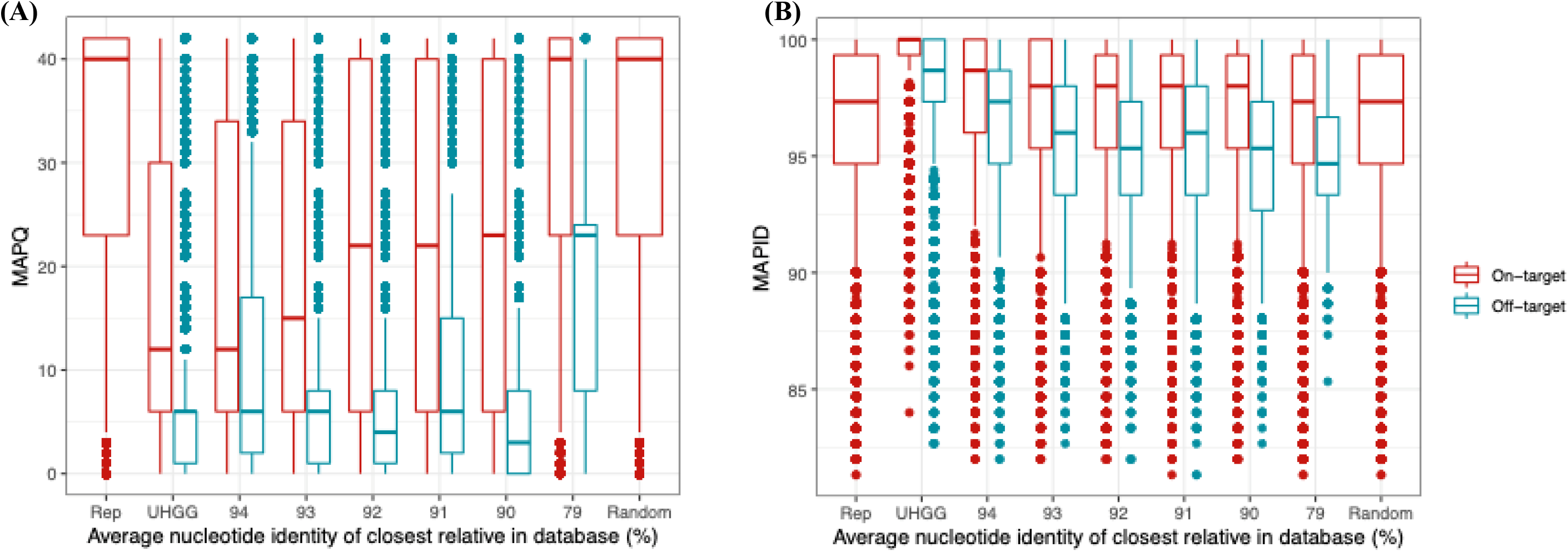
Closely related species greatly reduce alignment uniqueness. Simulated reads from *Collinsella sp003458415* were aligned to databases containing only the representative genome of the on-target species (Rep), two representative genomes: one on-target and from an one off-target species (horizontal axis: inter-species ANI), or representative genomes for 3,956 species (UHGG). No post-alignment filtering was applied. (**A**) A wide distribution of MAPQ values (vertical axis) with the Rep database shows that within-species multimapping reduces alignment uniqueness. Adding one closely related species to the database greatly reduces alignment uniqueness for reads aligned correctly to the on-target genome (red). This effect is correlated with inter-species ANI but remains fairly high out to 90% inter-species ANI. Using MAPQ > 30 for post-alignment filtering would remove the majority of on-target alignments. Including all 3,956 species in the database does not lead to a further decrease in MAPQ compared to including a closely related species with 94% inter-species ANI, emphasizing that low alignment uniqueness is mostly driven by highly related genomes in the database. Cross-mapped reads aligned to off-target genomes (blue) tend to have even lower MAPQ than on-target alignments. (**B**) As expected, reads aligned to the on-target genome tend to have high sequence identity (vertical axis: MAPID). This is especially true when there is a closely related species in the database (UHGG and 94% inter-species ANI), because many reads that would have lower MAPID are not correctly not aligned. Reads aligned to the off-target genome have slightly lower MAPID than on-target alignments, and their MAPID decreases with inter-species ANI.

Adding another species to the database drastically changes the distribution of MAPQ scores. For reads correctly aligned to the on-target genome, MAPQ drops precipitously when a closely related species (94% ANI) is added. Median MAPQ slowly increases as the inter-species ANI decreases, returning to a distribution similar to that with only the on-target genome when ANI < 80%. The distribution of MAPQ values for reads aligned to the off-target genome (cross-mapping) is lower than that of the on-target alignments across ANI values with the majority of alignments having MAPQ < 10 even at 94% ANI. In contrast, MAPID for reads correctly aligned to the on-target genome remains high with a closely related species in the database. While MAPID for cross-mapped reads tends to be lower than that of on-target reads, these distributions are highly overlapping.

Next, we repeated these analyses using a large, diverse reference database containing representative genomes for 3,956 species from the UHGG genome collection. With this many off-target species alignment uniqueness is even lower, with only 23.3% of the on-target reads aligning and a distribution of MAPQ values similar to when using only the species with 94% ANI to the on-target species. Meanwhile, MAPID for both on-target and off-target alignments is higher than any other simulation scenario, because only reads that are perfect or near-perfect matches are aligned. Altogether, these results make it very clear that incorrect alignments may have high scores and that closely related species severely reduce alignment uniqueness. Thus, while some form of post-alignment filtering is essential in the setting of metagenotyping in order to limit errors due to cross-mapping, a MAPQ threshold that works well in the absence of closely related species (e.g., MAPQ ≥ 30) will remove many correctly aligned reads.

To demonstrate this tension, we tracked the fate of every *C. sp003458415* read from the above analysis using the commonly employed post-alignment filter MAPQ ≥ 30 versus MAPQ ≥ 10. We chose MAPQ ≥ 10 as an alternative post-alignment filtering threshold, because off-target alignments in **Figure 6A** tend to have MAPQ below 10, including when using the database with 3,956 UHGG species, while many on-target alignments have MAPQ between 10 and 30. In both cases, we used MAPID ≥ 94% to remove some off-target alignments while not filtering out too many on-target ones (**Figure 6B**). We classified each read based on whether it was unaligned, incorrectly aligned to the off-target species, or correctly aligned to the on-target species before filtering, plus whether it passed the MAPQ and MAPID filters or not. Across the series of databases where the off-target genome has varying similarity to the on-target genome (90-95% inter-species ANI), we observed that MAPQ ≥ 10 enables many more reads to be mapped to the on-target genome compared to using MAPQ ≥ 30 (**Figure S4**). Most of these reads pass post-alignment filtering when only the on-target genome is in the database. Meanwhile, cross-mapping is higher at MAPQ ≥ 10 versus 30, as expected, but this increase is relatively small. These results illustrate that a closely related genome in the database can have a bigger impact on alignment uniqueness than on cross-mapping and that retaining alignments with medium values of MAPQ could be advantageous.

## ADJUSTING POST-ALIGNMENT FILTERS INCREASES METAGENOTYPE RECALL WITHOUT LARGE NUMBERS OF FALSE POSITIVES

Post-alignment filtering is used in metagenotyping tools to remove reads whose alignments are not unique or high scoring enough to be confident that the aligner has mapped the read to the correct species and genomic location. The goal is to do this without removing reads that are aligned correctly. Our results above indicate that balancing these two objectives can be difficult when there are closely related species in the database that compete for reads.

To explore this dilemma, we conducted simulations to quantify the effects of various post-alignment filters on metagenotypes across species. For these experiments, we used 86 diverse bacterial species whose closest relatives in the genome database had a range of inter-species ANI values (78-95%). First, we computed the recall and precision of SNVs without using any post-alignment filtering compared to different choices of MAPQ-based filtering (**Figure 7A**). This analysis showed that precision tends to be very high overall but lower for alternative alleles (median = 99.95% compared 99.99% for reference alleles). Precision is negatively correlated with inter-species ANI regardless of allele type, underscoring the importance of post-alignment filtering to avoid false positive SNVs. As expected from the rates of multimapping and reduced alignment uniqueness we quantified above, SNV recall is affected by post-alignment filters. Recall drops as the MAPQ threshold is increased, due to the presence of second-best alignments with scores that are not a lot worse than the best alignment. Overall, recall is lower for alternative versus reference alleles, especially when the closest off-target genome has higher inter-species ANI.

**Figure 7.**
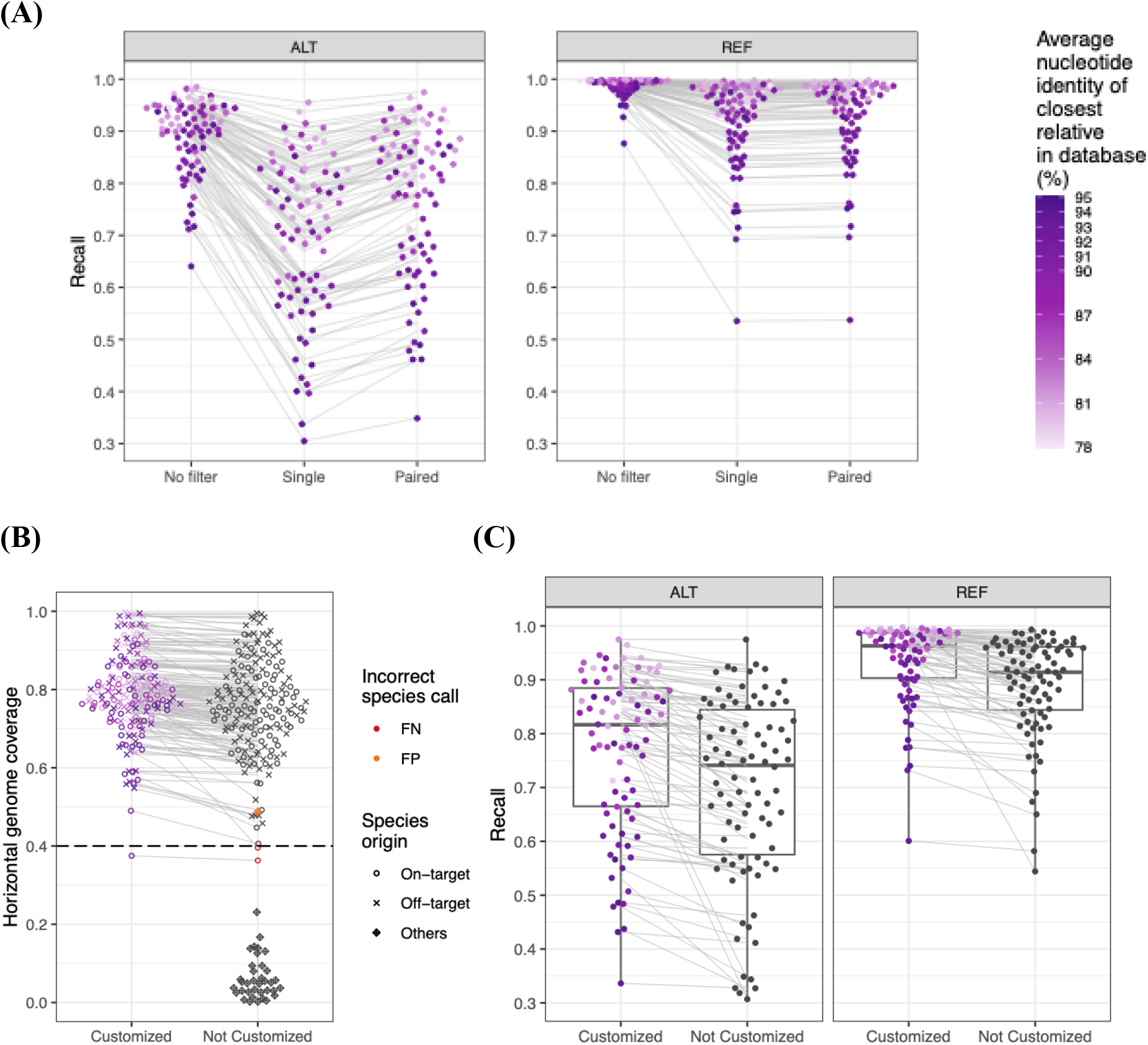
Post-alignment filtering and database customization improve SNV recall. To explore potential solutions to metagenotyping errors caused by closely related species in the database, we simulated reads from 86 species and aligned them to a database containing representative genomes for the on-target species and one off-target species with varying inter-species ANI (shade of purple). (**A**) SNV recall is reduced with closely related species (darker purple), especially for alternative (ALT versus REF) alleles. Recall falls further when individual reads are subjected to postalignment filtering (Single: MAPQ at least 10, MAPID at least 94), but increases when paired-end filtering is used (MAPQ at least 10 for at least one read in a proper pair), with 8% more SNVs correctly genotyped at ALT sites. (**B**) Adding reads from the off-target species to the simulated metagenomes, we found that a horizontal coverage threshold of 40% (dashed line) accurately distinguishes most species in the reads (on-target, off-target) from species not present (Others), with one false positive and two false negative species when the database is not customized (3,956 UHGG genomes). Customizing the database to include only the two species in the reads increases horizontal genome coverage, eliminating one false negative and the false positive species. (C) The customized database increases SNV recall at both ALT (7.5%) and REF (4.9%) sites.

Leveraging paired-end reads helps reduce this drop in recall without degrading precision. Specifically, we recommend to use only properly-aligned read pairs for metagenotyping (i.e., both ends of the reads are properly oriented and mapped within a reasonable distance given the expected distance input to the alignment software) and to retain both reads whenever one read has a sufficiently high MAPQ and MAPID. Across all 86 species, paired-end filtering with MAPQ ≥ 10 and MAPID ≥ 94% increases recall to values intermediate between no filter and filtering each read independently. The paired-end filter is particularly helpful for alternative alleles. It works for two reasons: (1) a read with low sequence similarity can be rescued if its pair has very high MAPID, and (2) a read with low alignment uniqueness (MAPQ < 10) can be rescued if its pair has a unique alignment. With the Bowtie2 aligner, it is important to properly set up the -X (maximum fragment length) to guarantee read pairs from DNA fragments longer than 500 nucleotides will be aligned (Zhao et al., 2022). Requiring proper pairs does filter out some improperly-aligned individual reads, so the total number of reads passing post-alignment filtering is similar for paired-end and single-end filtering. We conclude that choosing a MAPQ threshold appropriate to the community being studied and applying paired-end filtering together provide a good balance between false positive and false negative metagenotypes.

## DATABASE CUSTOMIZATION TO SAMPLE

Next, we considered horizontal coverage as a way to avoid false positive species and vertical coverage as a way to identify local regions affected by cross-mapping. These coverage filters can be applied after post-alignment filtering and pileup, just before calling SNVs. Our simulations showed that both of these commonly accepted quality measures can be as high for closely related off-target genomes as they are for the on-target genome, making them poor choices for eliminating erroneous alignments and reducing metagenotype error. However, we observed that horizontal coverage > 40% works well for determining if a species is present in the metagenome, resulting in only 1/86 false negative species in our simulation (**Figure 7B**). Being able to determine which species are in fact present motivated us to examine customization of the database to the metagenomic sample as a way to mitigate the effects of closely related species. Some strain-level analysis pipelines have incorporated a species detection or taxonomic prescreening step to customize the database to present species. For example, MIDAS2 (Zhao et al., 2022) uses the median coverage of 15 universal single copy genes, while HUMAnN2 (Franzosa et al., 2018) uses MetaPhlAn2 (Truong et al., 2015) to rapidly identify community species.

To test if adding genomes not present in the sample to the database can reduce metagenotype accuracy, we conducted a simulation in which reads from 86 pairs of closely related species (inter-species ANI > 92%) were combined in a metagenomic sample and then alignment, paired-end filtering, and SNV calling were performed with two database options: (i) a database that only includes the two species in the sample (customized) versus (ii) a database that includes 3,956 UHGG species (not customized). As expected, we noted that alignment was faster with the smaller, customized databases, although this speed up was canceled out by the computing resources needed for determining which species were present in the metagenome. Next, we looked at horizontal genome coverage and observed that the genomes for both species in the sample tend to have slightly higher coverage with the customized database (**Figure 7B**), and coverage is higher for metagenomes where the two species have low inter-species ANI (consistent with **Figure 4A**). For *Lachnospira eligens* (GCF_020735745.1, inter-species ANI: 94%, intra-species ANI: 95.6%), alignment uniqueness is so low that horizontal coverage is below a detection threshold of 40% with both databases despite being in the metagenome, while *Adlercreutzia equolifaciens* (GCF_000478885.1, inter-species ANI: 80.8%, intra-species ANI 95.9%) is correctly detected at this threshold with the customized database only. In general, UHGG species not in the sample have horizontal coverage below 40%, although uncultured *Adlercreutzia sp*. (MGYG-HGUT-02712) is falsely detected at this threshold and would be genotyped incorrectly by most pipelines. Importantly, we found that customized databases notably improved our power to detect SNVs, increasing recall 7.5% for the alternative allele and 4.9% for the reference allele (**Figure 7C**). Recall is higher for reference versus alternative alleles and for species pairs with low inter-species ANI (consistent with **Figure 5**). Finally, metagenotype precision is similar for both databases. Together, these results offer support for the idea of trimming genomes from metagenotyping databases and leaving only those from species detected in the sample.

We also evaluated the choice of representative genome for species that are present in the metagenomic sample. Our results and prior work (Olm et al., 2021) suggest that when a species has multiple genome sequences, selecting a reference genome that is as similar as possible to the genome in the metagenomic sample will increase alignment uniqueness and reduce mapping errors (**Figure 5A**). How best to pick representative genomes is not a solved problem. For a single sample containing a single strain, recall of reference alleles is highest when using the representative genome closest to that strain. But recall of alternative alleles is not correlated with genome-wide nucleotide identity, most likely because the average for the genome is not predictive of what happens at the most divergent sites. Furthermore, metagenomes containing two or more divergent lineages of a species make the choice of representative genome harder, as does identifying a single best genome to use for a set of diverse samples. We examined one species, *E. hormaechei_A*, in detail and found that using a centroid of all sequenced genomes for the species is an acceptable compromise that improves alignment rate ~10% compared to using the most distantly related genome (**Figure S5**). Although underutilized in practice, customization of reference databases shows great promise for maximizing alignment rate and minimizing reference bias.

## CONCLUSIONS AND PERSPECTIVES

Not long ago, the major challenge for reference-based metagenotyping was a paucity of species with a high-quality genome assembly. Now in some environments and some taxonomic groups, we have in a sense too many genomes, or rather too many for the existing tools to work in the intended way. From the perspective of reference-based metagenotyping, database growth has pros and–perhaps surprisingly–also cons. In this Synthesis, we quantified these drawbacks and the effects of potential bioinformatics solutions, highlighting how databases containing closely related species reduce alignment uniqueness and increase metagenotype errors.

Integrating across all our results, we identified low alignment uniqueness as one of the most important factors influencing metagenotype accuracy. One might think that diverse genome databases are a good thing, because adding the genome of a new species could prevent erroneous alignments of reads from that species to the genomes of related species. While this does happen, we found that a much more common outcome when a close species’ genome is added to a metagenotyping database is that reads are not unique enough to be aligned, especially when post-alignment filters are applied (e.g., MAPQ > 30). Another set of problems arise when reads align better to an off-target genome. We showed that cross-mapping can affect a large proportion of the genome and is worse when the on-target reference genome is diverged from the strain in the metagenomic sample. Low alignment uniqueness and cross-mapping are both worse for reads carrying alternative alleles compared to the reference genome.

These findings point to several actionable solutions. First, we recommend lowering the MAPQ threshold. In our analyses of hundreds of species with a variety of intra-species and inter-species ANI values, MAPQ > 10 emerged as a reasonable trade-off between false positive and false negative metagenotypes. Stricter thresholds do increase the accuracy of allele calls at sites that are metagenotyped, but at the cost of lower recall especially for SNVs with alternative alleles. Second, leveraging paired-end reads helps to increase SNV recall, which is especially important for alternative alleles and species with a close neighbor in the database. Finally, database customization can help in two ways. By using only the genomes of species likely to be in the metagenomic sample, many closely related genomes can be eliminated from the database, thereby mitigating their negative effects on metagenotypes. For species that are present, additionally selecting a reference genome that is similar to the genome in the reads will reduce reference bias and decrease alignment competition with related species. Some of these options are already available in existing metagenotyping tools (**Table S1**). To enable users to tune each step of analysis to their application, we encourage further development of pipelines in which the metagenotyping methods are fully customizable, from reference database to post-alignment filtering and SNV calling.

Even if all of these recommendations are followed, metagenotypes may still have fairly high error rates in some situations. Looking ahead, we can envision several large changes that could further reduce these errors. Metagenotyping tools could start using recent innovations in alignment algorithms, such as graph-based aligners (Kim et al., 2019), probabilistic alignment of multi-mapping reads (Bray et al., 2016; Shah and Ruthenburg, 2021; Vainberg-Slutskin et al., 2022; Zheng et al., 2019), and methods that utilize multiple reference genomes (Chen et al., 2021b). These strategies could reduce reference bias and resolve some cases of cross-mapping, though they increase compute time and memory use. Benchmarking these methods in the context of metagenotyping (Andreu-Sanchez et al., 2021) would reveal if alignment uniqueness increases and/or cross-mapping decreases, as well as the computational resources needed to achieve performance advantages. Another way to increase SNV accuracy, while also disentangling strains of the same species present within the a sample, may be to metagenotype multiple co-occurring SNVs together using some combination of long reads (Chen et al., 2022; Xie et al., 2020; Yahara et al., 2021), haplotype assembly (Ghazi et al., 2022; Li et al., 2019; Pulido-Tamayo et al., 2015), and singlecell metagenomic sequencing (Cole et al., 2020). The idea is to leverage information from genetically linked sites to increase confidence in metagenotypes. Similar to how paired-end reads increase recall of SNVs, we expect that these emerging techniques could rescue reads that would otherwise be filtered out, in particular closing the gap in recall we detected between reference and alternative alleles. Finally, we remind readers that performance decreases as a function of species abundance in simulations with different simulated coverage values (**Figure S1**), indicating more work is needed for applying the strategies in this study to low abundance bacteria. Matched amplicon sequencing (e.g., 16S) or metatranscriptomic data (RNA rather than DNA) may help with detection of low abundance species and is an interesting future direction for database customization. While all these approaches will require new or significantly re-engineered metagenotyping pipelines, their benefits may justify this effort.

This Synthesis focuses on reference-based metagenotyping, where alignment errors are the major source of inaccurate results. But alternatives to read alignment exist (Shajii et al., 2016). Inspired by forensic and taxonomic profiling tools that use exact matching of short sequences (k-mers) (Breitwieser et al., 2018; Liu et al., 2019; Ounit et al., 2015; Phillippy et al., 2009), we developed a metagenotyping pipeline in which the database is comprised of k-mers covering each allele of known SNV sites, filtered to remove any k-mers that occur in any other sequenced genome (Shi et al., 2022). This approach reduces both false positive and false negative SNV calls, and it is faster than alignment. However, very few SNVs are covered with unique k-mers when there are closely related species in the database, which limits the number of SNVs that can be metagenotyped per species. Also, this strategy is specific to SNVs identified by comparing reference genomes, for which k-mers can be designed, and the k-mers must not contain flanking insertions or deletions that interfere with exact matching. Another emerging strategy is completely reference-free metagenotyping in which reads are directly compared to each other to detect SNVs, insertions and deletions (Arif et al., 2019; Laso-Jadart et al., 2020; Leggett and MacLean, 2014; Peterlongo et al., 2017). Further benchmarking this approach on metagenomic data and developing parallel methods for larger structural variants are important future directions.

While our analyses focused on SNVs in bacterial genomes, most of the points raised here apply to other types of variants and to different taxonomic groups. Indeed, all of the metagenotyping challenges associated with diverse and closely related species will affect those lineages of archaea, viruses, and eukaryotes where genomes are being densely sampled and assembled (Emerson et al., 2018; Gregory et al., 2020; Gregory et al., 2019; Massana and López-Escardó, 2022; Mukherjee et al., 2017; Nayfach et al., 2021). Beyond SNVs, alignment is used in several tools that metagenotype gene copy number variants and other structural variants (Greenblum et al., 2015; Zeevi et al., 2019; Zhao et al., 2022). Since these variants are called based on coverage in pileups, their detection and quantification will be biased by reads that are not aligned, fail post-alignment filtering, or are incorrectly aligned. Therefore, competition for reads across species, within-species multimapping, and cross-mapping will affect structural variant metagenotypes in ways qualitatively similar to the effects we demonstrated for SNVs. More broadly, our findings are also very relevant to metagenomic analyses that do not involve genotyping, such as species detection and abundance estimation, where probabilistic mapping has been recently proposed as a solution for perfectly multi-mapping reads (Vainberg-Slutskin et al., 2022). Thus, many aspects of microbiome bioinformatics require careful consideration of how alignment algorithms perform on a tree of life in which many lineages are now densely sequenced.

Despite the prevalence of closely related species in genome databases today, it is important to remember that the vast majority of species, including most bacteria, still have limited or no genome sequences. We emphasize that genome sequencing aimed at expanding reference genome collections should focus on capturing these under-represented lineages. However, this Synthesis shows that alignment and other bioinformatics tools must continue to evolve in order to remain accurate in the face of closely related genomes.

## STAR * METHODS

### Software and algorithms

fastANI (version 1.33) (Jain et al., 2018)

Mash (version 2.2) (Ondov et al., 2016)

ART (version 2.5.8) (Huang et al., 2012)

Bowtie2 (version 2.3.5.1) (Langmead and Salzberg, 2012)

MUMmer4 (version 4.0) (Marcais et al., 2018)

MIDAS2 (version 0.5) (Zhao et al., 2022)

R (version 4.2.0) (R Core Development Team, 2022) (packages: ggplot, ggbeeswarm, ggsci)

metacoder (version 0.3.5) (Foster et al., 2017)

### Databases

NCBI Assembly (May 29, 2022) (Kitts et al., 2015)

Genome Taxonomy Database (GTDB; version R207) (Parks et al., 2021)

Genomes of Earth’s Microbiomes (GEM; June 1, 2022) (Nayfach et al., 2021)

Unified Human Gastrointestinal Genome collection (UHGG; v1.0) (Almeida et al., 2021a)

### Survey of prokaryotic genome collections

We counted the number of prokaryotic species with a genome in the NCBI Assembly database from 1999 to 2022. NCBI Assembly was used for this analysis because (1) it is the largest genome database, (2) it is updated daily, and (3) it unambiguously identifies and tracks changes. To ensure maximum inclusiveness as well as low redundancy, we counted unique species of bacteria and archaea with any of four levels of genome assemblies: complete genome assemblies, assemblies that include chromosomes or linkage groups, scaffolds and contigs, assemblies that include scaffolds and contigs, and assemblies that include only contigs. We also surveyed the number of species and the number of genomes per species in the current versions of three other databases: GTDB, GEM, and UHGG. These large genome collections have distinct features. GTDB contains mainly assemblies from isolate whole-genome shotgun sequencing projects, and it uses genomes solely from the NCBI Assembly database. GEM is a collection of genomes from diverse environments. UHGG is a collection of genomes from the human gut microbiome. Both GEM and UHGG contain a high proportion of metagenome-assembled genomes (MAGs). Using pairs of high-quality GTDB genomes for *B. mallei* and *E. hormaechei_A*, we computed intra-species ANI. We aligned the metagenomic reads of 5 random US samples (ERR4330028, ERR4330046, ERR4334225, ERR4335281 and ERR4341723) and 5 random UK samples (ERR4334072, ERR4334226, ERR4335245, ERR4335298 and ERR4341720) from the PREDICT cohort (Asnicar et al., 2021) to UHGG (4,643 gut genomes) and NCBI (2013; 6,549 genomes) using Bowtie2 and computed the median alignment rates. These samples have very high sequencing depth (~55 million reads per sample on average), and none of them contributed genomes to UHGG or the 2013 release of NCBI.

### Survey of genomic similarity between species in GTDB

We downloaded a total of 65,703 genomes from GTDB and selected one high-quality representative genome for each of the 19,754 species. For each representative genome, we used Mash to generate a 21-mer sketch profile (*mash sketch -k 21 -s 5000*) and calculated pairwise genomic distance to all other representative genomes (*mash dist*). Mash distance estimates genome-wide average nucleotide identity (ANI) and is computationally feasible with 19,754 genomes. We denoted a species as having a closely related species (CRS) if the smallest Mash distance to any other species was below 0.08 (≥ *92%* ANI). The heat_tree function in Metacoder was used to visualize the number of species with CRS on the GTDB phylogeny, rendered as a cladogram.

### Quantification of intra-species and inter-species genomic similarity

To assess genomic diversity within species, we used intra-species ANI. For each species, we used fastANI to compute the sequence similarity between conspecific genome assemblies. This calculation was applied to all high-quality genomes of selected species from UHGG (1,969 species with 2-6,645 genomes per species) and to all high-quality genomes of selected species from GTDB (9 species with 1,000-2,000 genomes per species). We also used intra-species ANI to compare individual high-quality genomes from NCBI Assembly to conspecific genomes from UHGG. The resulting intra-species ANI values were used to assess how diverse the genomes from different species are and to select genomes for simulation experiments.

To assess genomic distance between species, we used inter-species ANI. For each pair of species, we used fastANI to compute the similarity between a representative genome of each species. This calculation was applied to high-quality representative genomes of species pairs from UHGG. The resulting inter-species ANI values were used to select genomes for simulation experiments.

### Metagenomic simulations

To evaluate alignment and metagenotyping errors across a broad range of scenarios, we simulated metagenomic sequencing reads from UHGG, NCBI Assembly, and GTDB genomes selected based on intra-species and inter-species ANI (**Table S2**). We used two GTDB species with >1,000 genomes (*Burkholderia mallei* and *Enterobacter hormaechei_A*) to evaluate reference bias (**Table S3**). We used 327 UHGG species with at least two high-quality genomes and at least one CRS (interspecies ANI ≥ 92%) to further explore reference bias and to evaluate cross-mapping, alignment uniqueness, and alignment sequence identity (**Table S4**). We used 86 NCBI genomes from species commonly found in the human gut (Cheng et al., 2021) and represented in UHGG with at least one related species (inter-species ANI ≥ 80%) to evaluate performance differences between reference and alternative alleles, as well as the effects of post-alignment filtering and database customization (**Table S5**). All genomes used as simulation templates were high quality (Completeness ≥ 90, Contamination ≤ 5), and only species with at least two high-quality genomes were used.

For each genome used as a simulation template, 150-basepair, paired-end Illumina sequencing reads were computationally generated using ART (GTDB and UHGG genomes: *art_illumina -ss HS25 -l 150 - m 1000 -s 100 -sp*, NCBI genomes: *art_illumina -ss HS25 -l 125 - m 600 -s 60 -sp*) at a range of genome coverage levels (1X - 50X). In an initial set of experiments, only one species was included in the metagenomic reads. Template genomes were selected to have a range of values for intra-species ANI to the representative genome of that species in the metagenotyping database (see below). Next, we evaluated the effects of adding reads from additional species with varying inter-species ANI to the first species (based on representative genomes). Since the majority of errors that we detected were due to the most closely related species in the metagenomic reads, we simplified further experiments by simulating reads from only two species, one designated as the on-target species and the second as the off-target species.

### Metagenotyping databases and read alignment

For each iteration of the simulation experiments, metagenomic reads generated from the template genome(s) were aligned to a particular database using Bowtie2 (*bowtie2 --no-unal -X 1000.0 --end-to-end --very-sensitive*). In order to tune ANI between the reads and the reference database, we selected representative genomes based on intra-species and inter-species ANI (**Table S2**). To evaluate reference bias, reads simulated from GTDB and UHGG non-representative genomes were aligned to the default representative genome for their species. GTDB reads were also aligned to the centroid of all GTDB genomes (lowest average pairwise ANI) and a boundary genome (highest average pairwise ANI). Reads simulated from NCBI genomes were aligned to a UHGG genome with ANI 95.5% - 99.5% to the NCBI genome (either the default representative genome or another genome if the default one is outside this ANI range). For all other analyses, we used databases with (i) only the UHGG representative genome of the on-target species, (ii) the UHGG representative genomes of the on-target and off-target species, or (iii) all UHGG representative genomes.

### Defining ground truth variants with whole-genome alignment

To determine the correct genotypes for simulated metagenomic reads, we compared the template genome to the representative genome in the metagenotyping database. Whole-genome alignments of pairs of conspecific genomes were aligned using *nucmer* in the MUMmer package (*--mum*: only use anchor matches that are unique in both the reference and query). Poorly aligned blocks with average sequence identity < 95% were identified, and nucleotides in these blocks were excluded from metagenotype performance assessments. For retained blocks, single nucleotide variants (SNVs) were called and used as ground truth variant sites for simulation experiments. All matching sites in these blocks were used as ground truth non-variant sites.

### Metagenotype analysis

We used MIDAS2 to metagenotype each simulated metagenome with each choice of database. Single-sample metagenotyping was performed using the SNV module of MIDAS2 (*midas2 run_snps*). For each sample, MIDAS2 reports summary statistics of the read alignment and pileup. SNVs were called in the pileup of metagenomic reads, and the persample per-species major allele per-site was compared to ground truth variant and non-variant sites from whole-genome alignments (see above). MIDAS2 is flexible enough that we could explore horizontal and vertical coverage thresholds, post-alignment filters, and database customization across a range of settings that cover most of the defaults in other metagenotyping tools (**Figure S1**). Performance differences between MIDAS2, metaSNV2, and inStrain have been investigated elsewhere (Olm et al., 2021; Zhao et al., 2022).

### Post-alignment filtering

In an initial set of experiments, we assessed alignment and metagenotyping errors without using any post-alignment filters. Then these performance results were compared to results with post-alignment filtering. Three different post-alignment filters were implemented by customizing the MIDAS2 command line: no filter (*--analysis_ready*), single-end based filter (default option), and paired-end filter (*--paired_only*).

### Performance assessments

We evaluated performance using definitions that adhere to best practices for microbial genomics (Olson et al., 2015). Alignment rate was calculated as the number of aligned reads (after any filter was applied) divided by the total number of reads. MIDAS2 computes horizontal coverage as the proportion of genomic sites aligned with at least two post-filtered reads, and vertical coverage as the average read depth of genomic sites covered with at least two reads. SNV precision was computed as the number of correctly called non-variant or variant sites over the number of called non-variant or variant sites. SNV recall was computed as the number of correctly called non-variant or variant sites over the total number of nonvariant or variant sites in the ground truth sets defined from whole-genome alignments. Precision provides insight into accuracy of the metagenotyping results, while recall measures statistical power.

## SUPPLEMENTARY INFORMATION

## ACKNOWLEDGEMENTS

This work was funded by NSF grant #1563159, Chan Zuckerberg Biohub, and Gladstone Institutes.

## AUTHOR CONTRIBUTIONS

CZ and KSP conceived the project and simulation methodology. CZ and ZJS wrote code, designed and performed computational experiments, and made figures. CZ, ZJS, and KSP made tables. All authors interpreted the results. KSP drafted the manuscript. CZ and ZJS edited the manuscript.

## Supplemental Figures with Captions

**Figure S1.**
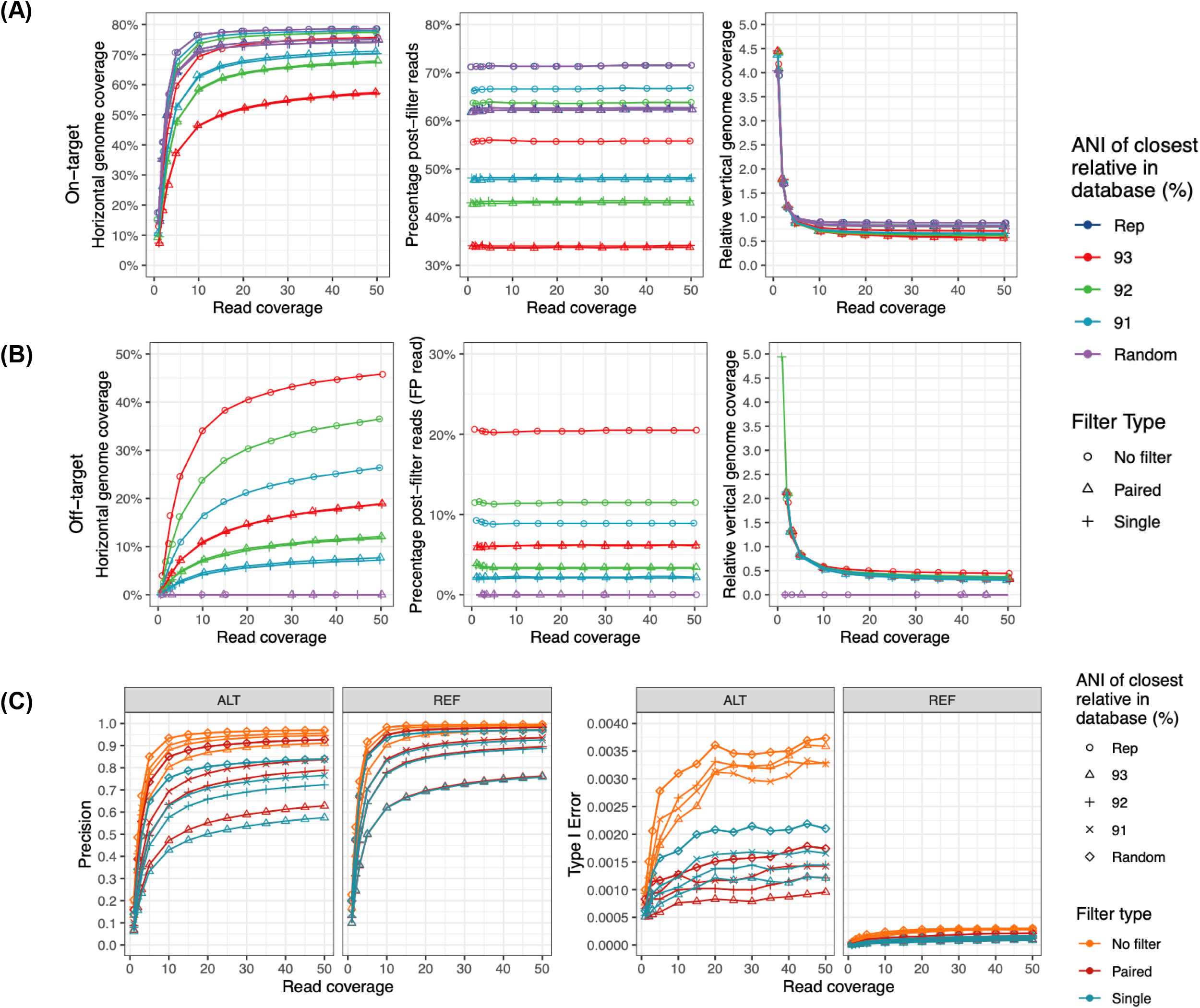
Convergence of performance statistics around 20x coverage. Reads simulated from *Catenibacterium mitsuokai DSM-15897* (GCF_000173795.1) with coverage ranging from 1X to 50X (horizontal axes) were aligned to a database containing the UHGG representative genomes for *C. mitsuokai* (On-target) and another species (Off-target) selected to have a range of interspecies ANI (Rep: only the on-target genome, 91-93: ANI between representative genomes of two species in the database, Random: inter-species ANI < 77). Post-alignment filtering was varied (No filter, Single-end filter, Paired-end filter) with MAPQ at least 10 and MAPID at least 94 as thresholds. (**A-B**) For each representative genome in the database, we tracked the following alignment statistics after any post-alignment filtering and dropping sites with only one aligned read: percentage of genome covered by aligned reads (horizontal genome coverage; left), percentage of reads aligned to that genome (middle), and average per site read depth in the pileup divided by the simulated coverage (relative vertical genome coverage; right). (**A**) Alignment statistics for the on-target species *C. mitsuokai DSM-15897* are lower with a closely related species in the database, with post-alignment filtering, and at simulated coverage values below 10X. Even when the database contains only the on-target representative genome and no postalignment filtering is performed, horizontal coverage is only ~80% and ~28% of reads are not aligned due to sequencing errors, reference bias, and other factors that prevent accurate read mapping. (**B**) Alignment statistics for the off-target genome show that cross-mapping is prevalent. It increases with simulated coverage, but this relationship levels off around 10X-20X coverage depending on the alignment statistic. Cross-mapping depends heavily on inter-species ANI and is mitigated by post-alignment filtering, illustrating the trade-off between maximizing on-target alignments while minimizing off-target alignments. (**C**) Metagenotyping was performed with MIDAS2 using the alignments from **A-B**, and SNV calls were compared to the gold standard of SNVs from whole-genome alignments of the simulation template genome of the on-target species to the representative genome of the on-target species. The majority allele was used for this evaluation, and ambiguous sites (tied reads for both alleles) were excluded. SNV precision (correctly called sites divided by all unambiguous sites with at least 2 reads; left) is lower with post-alignment filtering, lower simulated coverage, and a closely related species in the database. It is lower for alternative (ALT) versus reference (REF) alleles in most scenarios. SNV type I error (incorrectly called sites divided by all unambiguous sites with at least 2 reads; right) is high overall, but shows similar trends. It is notably lower for ALT alleles.

**Figure S2.**
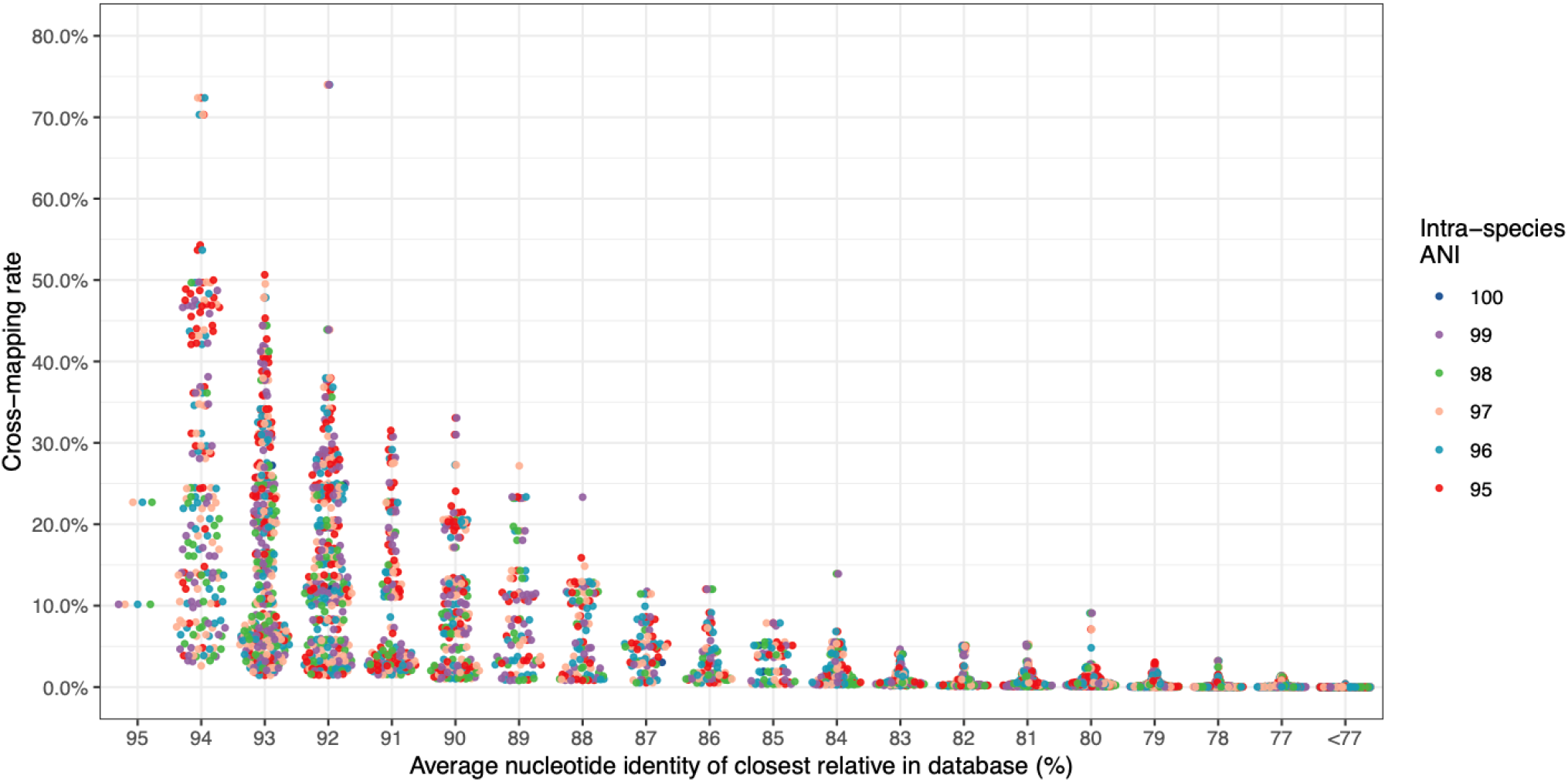
Cross-mapping is mutual. The simulation in **Figure 5** was augmented by including reads from a genome from the off-target species. The metagenome containing reads from both species was aligned to the same database containing the representative genomes for both species. No post-alignment filtering was applied. Mirroring the result in **Figure 5A**, the cross-mapping rate for reads from the off-target species to the on-target representative genome was high and positively correlated with inter-species ANI (horizontal axis). As expected, it was not highly correlated with the intra-species ANI of the on-target species (colors).

**Figure S3.**
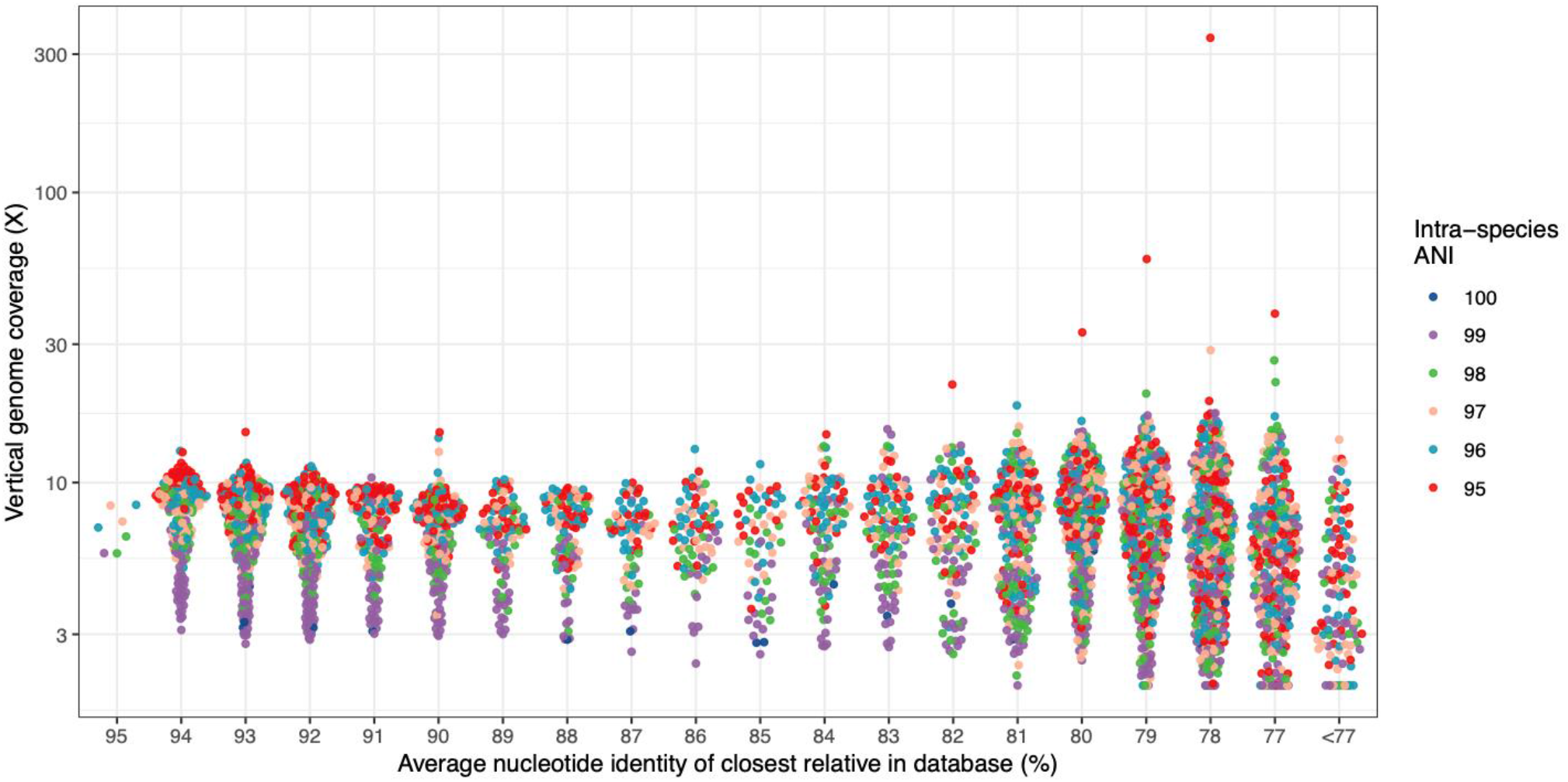
Cross-mapping leads to high vertical genome coverage. Using the simulations from Figure 5, we observed that an average site in the off-target genome can have many reads aligned. The reads were simulated at 20X coverage, so 10X vertical coverage represents half of the expected coverage for the on-target genome. Vertical coverage is higher when the simulation template genome has lower similarity to the on-target representative genome, but it does not show a strong trend with inter-species ANI. Vertical coverage is the mean read depth in the pileup, counting only crossmapped reads.

**Figure S4.**
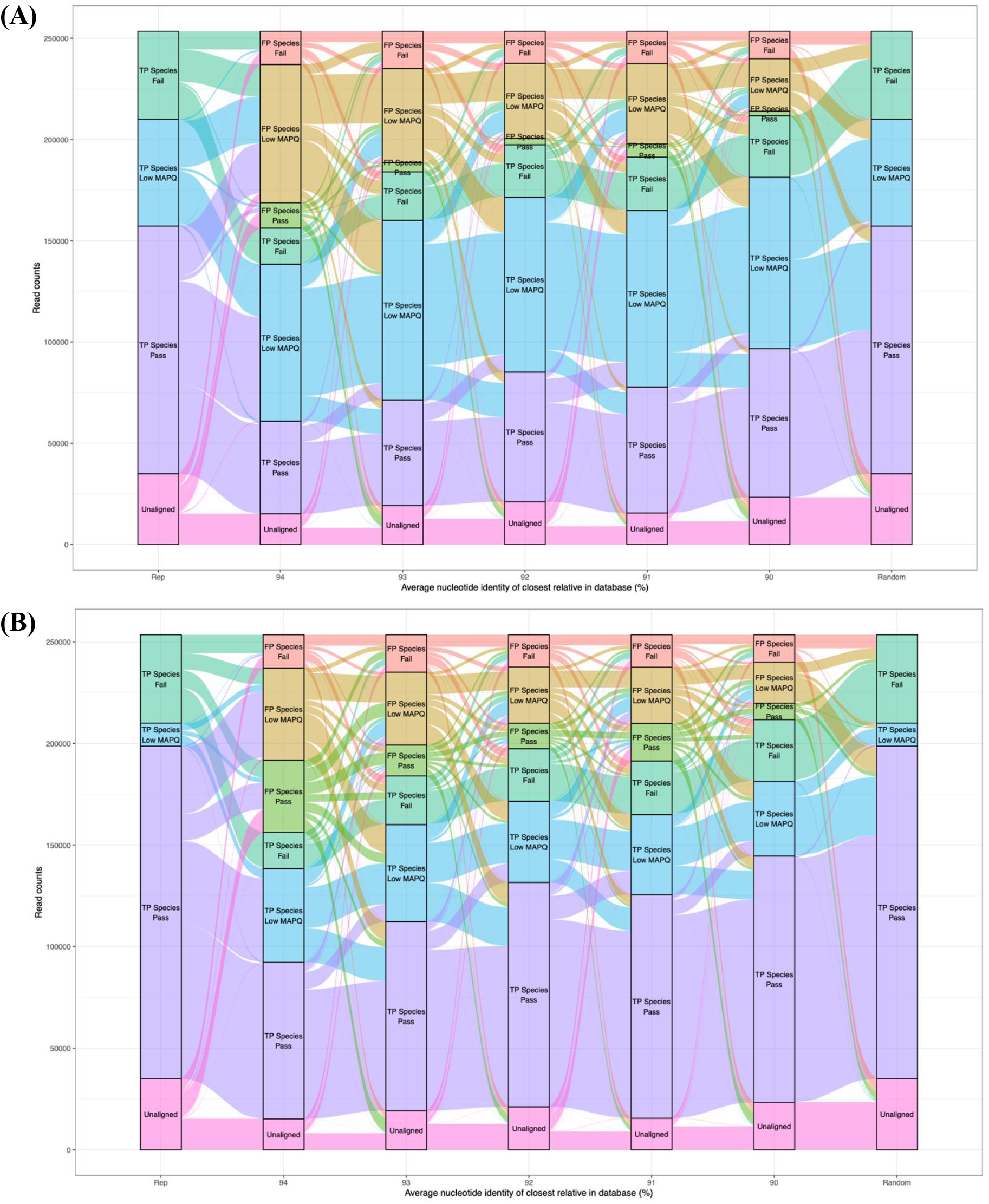
Tracking the fate of every read as a function of database and post-alignment filters. We simulated reads from *Collinsella sp003458415* (UHGG species ID: 100197, intra-species ANI ~95%) at 20X coverage and aligned them to databases containing representative genomes for *C. sp003458415* (on-target) and another species (off-target) selected to have a range of inter-species ANI (Rep: only the on-target genome, 90-94: ANI between representative genomes of two species in the database, Random: a randomly selected UHGG species). Post-alignment filtering was performed with a MAPID threshold of 94% and a MAPQ threshold of 10 versus 30. Reads are traced across databases and classified in each analysis as follows: Unaligned (pink), TP Species Pass (correctly aligned reads with MAPID and MAPQ above thresholds; purple), TP Species Low MAPQ (aligned to on-target genome with MAPID at least 94% but MAPQ below threshold; blue), TP Species Fail (aligned to on-target genome with MAPID and MAPQ below threshold; teal), FP Species Pass (aligned to off-target genome with MAPID and MAPQ above thresholds, i.e., cross-mapped; lime), FP Species Low MAPQ (aligned to off-target genome with MAPID at least 94% but MAPQ below threshold; orange), FP Species Fail (aligned to off-target genome with MAPID and MAPQ below thresholds; red). (**A**) At a MAPQ threshold of 30 (common default), closely related species cause cross-mapping (lime) and greatly reduce alignment uniqueness (purple reads flowing to blue and orange). (**B**) At a MAPQ threshold of 10 more correctly mapped reads pass post-alignment filtering (purple) at the cost of more cross-mapping (lime).

**Figure S5.**
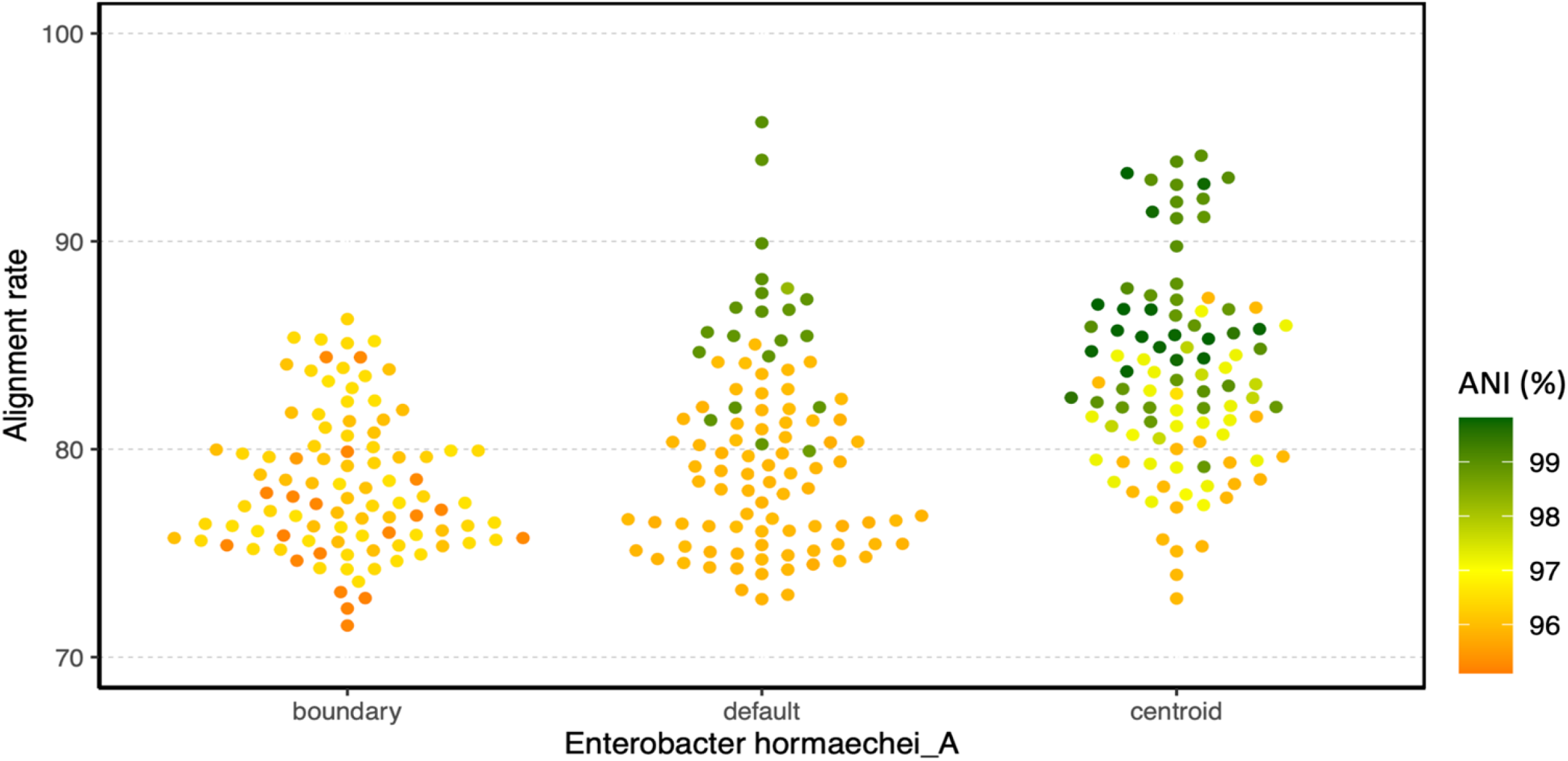
Reference bias as a function of representative genome. We simulated reads from 100 diverse GTDB genomes of *Enterobacter hormaechei_A* at 30X coverage and aligned them to databases containing three different choices of on-target reference genome: boundary (furthest from other GTDB *E. hormaechei_A* genomes, mean intra-species ANI = 96.0%; GCF_008082005.1), default GTDB representative (mean intra-species ANI = 96.6%; GCF_001729745.1), or centroid (closest to other GTDB *E. hormaechei_A* genomes, mean intra-species ANI = 97.7%; GCF_003964925.1). For each reference genome, alignment rate decreases with ANI to the simulation template genome (color scale). Across reference genomes, alignment rate tends to be highest for the centroid, followed by the GTDB default representative and then the boundary genome.

**Table S1.** Comparison of features implemented in different alignment-based metagenotyping tools.

| Metagenotyping pipeline | Reads aligned to | Database customization | Post-alignment filter uses paired-end information | Default MAPQ filter |
| --- | --- | --- | --- | --- |
| MIDAS2<br>(Zhao et al. 2022) | Whole genomes | Species detected by profiling 15 universal single copy genes | Yes | MAPQ >= 10 |
| inStrain<br>(Olm et al. 2021) | Whole genomes | No | Yes | MAPQ >= 2 |
| metaSNV v2<br>(Van Rossum et al. 2021) | Whole genomes | No | No | Uniquely aligned only. |
| StrainPhlan3<br>(Beghini et al. 2021) | Species-specific marker genes | No | No | MAPQ > 30 |

**Table S2.** Data and analysis used in metagenomic simulation figures.

| Figure | Dataset | Genome(s) used to simulate metagenomic reads in each iteration | Database | Filter |
| --- | --- | --- | --- | --- |
| Figure 3G | *B. mallei* | One *Burkholderia mallei* GTDB genome with varying ANI to the GTDB representative genome | GTDB representative genome for *B. mallei* (GCF_000011705.1) | None |
| Figure 3H | *E. hormaechei* | One *Enterobacter hormaechei*_A GTDB genome with varying ANI to the GTDB representative genome | GTDB representative genome for *E. hormaechei*_A (GCF_001729745.1) | None |
| Figure S5 | *E. hormaechei* | One *Enterobacter hormaechei*_A GTDB genome with varying ANI to the representative genome (boundary or centroid) | GTDB centroid genome for *E. hormaechei*_A (GCF_900077895.1) or GTDB boundary genome for *E. hormaechei*_A (GCF_008082005.1) | None |
| Figure S1 | *C. mitsuokai* | NCBI genome of *Catenibacterium mitsuokai* DSM-15897 (GCF_000173795.1) | UHGG representative genomes for *C. mitsuikai* <u>and</u> for one off-target species with varying ANI to the *C. mitsuikai* representative genome | None, Single, Paired |
| Figure 6A & 6B + Figure S4 | *C. sp003458415* | UHGG representative genome of *Collinsella sp003458415* (species ID: 100197) | UHGG representative genome for *C. sp003458415* <u>and</u> UHGG representative genome for one off-target species with varying ANI to the *C. sp003458415* representative genome <u>or</u> UHGG representative genomes for 3,956 species including *C. sp003458415* | None |
| Figure 4A | 327 UHGG species | One UHGG genome from the on-target species with varying ANI to the UHGG representative genome of that species | UHGG representative genome for on-target species | None |
| Figure 5A | 327 UHGG species | One UHGG genome from the on-target species with varying ANI to the UHGG representative genome of that species, one per 1% ANI bin (95% to 100%, up to five per on-target species) | UHGG representative genome for on-target species <u>and</u> UHGG representative genome for off-target species | None |
| Figure S2 | 327 UHGG species | One UHGG genome from the on-target species with varying ANI to the UHGG representative genome of that species <u>and</u> one UHGG non-representative genome from an off-target species with varying ANI to the on-target species, one per 1% ANI bin (77% to 95% based on the two representative genomes plus one random species, up to 19 per on-target species) | UHGG representative genome for on-target species <u>and</u> UHGG representative genome for off-target species |  |
| Figure 5B + Figure S3 | 327 UHGG species | One UHGG genome from the on-target species with varying ANI to the UHGG representative genome of that species<br>* Horizontal coverage only includes reads from the off-target species (cross-mapping) | UHGG representative genome for on-target species <u>and</u> UHGG representative genome for off-target species | None |
| Figure 4B & 4C | 86 NCBI strains | One NCBI genome from the on-target species | UHGG representative genome for on-target species <u>or</u> another UHGG genome for that species with ANI 95.5% - 99.5% to NCBI genome | None |
| Figure 7A | 86 NCBI strains | One NCBI genome for the on-target species | One UHGG representative genome <u>or</u> another UHGG genome for on-target species with ANI 95.5% - 99.5% to NCBI genome <u>and</u> one UHGG representative genome for the off-target species with varying ANI to the on-target species | None, Single, Paired |
| Figure 7B & 7C | 86 NCBI strains | One NCBI genome from the on-target species <u>and</u> one UHGG genome from the closest off-target species (based on ANI to the on-target species) | Customized: One UHGG representative genome <u>or</u> another UHGG genome for on-target species with ANI 95.5% - 99.5% to NCBI genome <u>and</u> one UHGG representative genome for the off-target species. Not customized: 3,956 UHGG representative genomes | Paired |

\* The genome used to simulate reads is never the same with the representative genome in the database.

## REFERENCES

Almeida, A., Nayfach, S., Boland, M., Strozzi, F., Beracochea, M., Shi, Z.J., Pollard, K.S., Sakharova, E., Parks, D.H., Hugenholtz, P., et al. (2021a). A unified catalog of 204,938 reference genomes from the human gut microbiome. Nature Biotechnology 39, 105–114.

Almeida, A., Nayfach, S., Boland, M., Strozzi, F., Beracochea, M., Shi, Z.J., Pollard, K.S., Sakharova, E., Parks, D.H., Hugenholtz, P., et al. (2021b). A unified catalog of 204,938 reference genomes from the human gut microbiome. Nat Biotechnol 39, 105–114.

Andreu-Sanchez, S., Chen, L., Wang, D., Augustijn, H.E., Zhernakova, A., and Fu, J. (2021). A Benchmark of Genetic Variant Calling Pipelines Using Metagenomic Short-Read Sequencing. Front Genet 12, 648229.

Anyansi, C., Straub, T.J., Manson, A.L., Earl, A.M., and Abeel, T. (2020). Computational Methods for Strain-Level Microbial Detection in Colony and Metagenome Sequencing Data. Front Microbiol 11, 1925.

Arif, M., Gauthier, J., Sugier, K., Iudicone, D., Jaillon, O., Wincker, P., Peterlongo, P., and Madoui, M.A. (2019). Discovering millions of plankton genomic markers from the Atlantic Ocean and the Mediterranean Sea. Mol Ecol Resour 19, 526–535.

Asnicar, F., Berry, S.E., Valdes, A.M., Nguyen, L.H., Piccinno, G., Drew, D.A., Leeming, E., Gibson, R., Le Roy, C., Khatib, H.A., et al. (2021). Microbiome connections with host metabolism and habitual diet from 1,098 deeply phenotyped individuals. Nat Med 27, 321–332.

Bray, N.L., Pimentel, H., Melsted, P., and Pachter, L. (2016). Near-optimal probabilistic RNA-seq quantification. Nat Biotechnol 34, 525–527.

Breitwieser, F.P., Baker, D.N., and Salzberg, S.L. (2018). KrakenUniq: confident and fast metagenomics classification using unique k-mer counts. Genome Biol 19, 198.

Bush, S.J., Foster, D., Eyre, D.W., Clark, E.L., De Maio, N., Shaw, L.P., Stoesser, N., Peto, T.E.A., Crook, D.W., and Walker, A.S. (2020). Genomic diversity affects the accuracy of bacterial single-nucleotide polymorphism-calling pipelines. Gigascience 9.

Chattopadhyay, S., Weissman, S.J., Minin, V.N., Russo, T.A., Dykhuizen, D.E., and Sokurenko, E.V. (2009). High frequency of hotspot mutations in core genes of Escherichia coli due to short-term positive selection. Proc Natl Acad Sci U S A 106, 12412–12417.

Chen, I.A., Chu, K., Palaniappan, K., Ratner, A., Huang, J., Huntemann, M., Hajek, P., Ritter, S., Varghese, N., Seshadri, R., et al. (2021a). The IMG/M data management and analysis system v.6.0: new tools and advanced capabilities. Nucleic Acids Res 49, D751–D763.

Chen, L., Zhao, N., Cao, J., Liu, X., Xu, J., Ma, Y., Yu, Y., Zhang, X., Zhang, W., Guan, X., et al. (2022). Short-and long-read metagenomics expand individualized structural variations in gut microbiomes. Nat Commun 13, 3175.

Chen, N.C., Solomon, B., Mun, T., Iyer, S., and Langmead, B. (2021b). Reference flow: reducing reference bias using multiple population genomes. Genome Biol 22, 8.

Cheng, A.G., Aranda-Díaz, A., Jain, S., Yu, F., Iakiviak, M., Meng, X., Weakley, A., Patil, A., Shiver, A.L., Deutschbauer, A., et al. (2021). Systematic dissection of a complex gut bacterial community. bioRxiv, 2021.2006.2015.448618.

Cole, B.J., Basso, J.T.R., and Visel, A. (2020). Power in isolation: insights from single cells. Nat Rev Microbiol 18, 364.

Deschamps-Francoeur, G., Simoneau, J., and Scott, M.S. (2020). Handling multi-mapped reads in RNA-seq. Comput Struct Biotechnol J 18, 1569–1576.

Emerson, J.B., Roux, S., Brum, J.R., Bolduc, B., Woodcroft, B.J., Jang, H.B., Singleton, C.M., Solden, L.M., Naas, A.E., Boyd, J.A., et al. (2018). Host-linked soil viral ecology along a permafrost thaw gradient. Nat Microbiol 3, 870–880.

Forbes, J.D., Bernstein, C.N., Tremlett, H., Van Domselaar, G., and Knox, N.C. (2018). A Fungal World: Could the Gut Mycobiome Be Involved in Neurological Disease? Front Microbiol 9, 3249.

Foster, Z.S., Sharpton, T.J., and Grunwald, N.J. (2017). Metacoder: An R package for visualization and manipulation of community taxonomic diversity data. PLoS Comput Biol 13, e1005404.

Franzosa, E.A., McIver, L.J., Rahnavard, G., Thompson, L.R., Schirmer, M., Weingart, G., Lipson, K.S., Knight, R., Caporaso, J.G., Segata, N., et al. (2018). Species-level functional profiling of metagenomes and metatranscriptomes. Nat Methods 15, 962–968.

Garrison, E., Siren, J., Novak, A.M., Hickey, G., Eizenga, J.M., Dawson, E.T., Jones, W., Garg, S., Markello, C., Lin, M.F., et al. (2018). Variation graph toolkit improves read mapping by representing genetic variation in the reference. Nat Biotechnol 36, 875–879.

Garud, N.R., and Pollard, K.S. (2020). Population Genetics in the Human Microbiome. Trends Genet 36, 53–67.

Ghazi, A.R., Munch, P.C., Chen, D., Jensen, J., and Huttenhower, C. (2022). Strain Identification and Quantitative Analysis in Microbial Communities. J Mol Biol, 167582.

Greenblum, S., Carr, R., and Borenstein, E. (2015). Extensive strain-level copy-number variation across human gut microbiome species. Cell 160, 583–594.

Gregory, A.C., Zablocki, O., Zayed, A.A., Howell, A., Bolduc, B., and Sullivan, M.B. (2020). The Gut Virome Database Reveals Age-Dependent Patterns of Virome Diversity in the Human Gut. Cell Host Microbe 28, 724–740 e728.

Gregory, A.C., Zayed, A.A., Conceicao-Neto, N., Temperton, B., Bolduc, B., Alberti, A., Ardyna, M., Arkhipova, K., Carmichael, M., Cruaud, C., et al. (2019). Marine DNA Viral Macro- and Microdiversity from Pole to Pole. Cell 177, 1109–1123 e1114.

Groussin, M., Poyet, M., Sistiaga, A., Kearney, S.M., Moniz, K., Noel, M., Hooker, J., Gibbons, S.M., Segurel, L., Froment, A., et al. (2021). Elevated rates of horizontal gene transfer in the industrialized human microbiome. Cell 184, 2053–2067 e2018.

Gunther, T., and Nettelblad, C. (2019). The presence and impact of reference bias on population genomic studies of prehistoric human populations. PLoS Genet 15, e1008302.

Hovhannisyan, H., Hafez, A., Llorens, C., and Gabaldon, T. (2020). CROSSMAPPER: estimating cross-mapping rates and optimizing experimental design in multi-species sequencing studies. Bioinformatics 36, 925–927.

Huang, W., Li, L., Myers, J.R., and Marth, G.T. (2012). ART: a next-generation sequencing read simulator. Bioinformatics 28, 593–594.

Jain, C., Rodriguez, R.L., Phillippy, A.M., Konstantinidis, K.T., and Aluru, S. (2018). High throughput ANI analysis of 90K prokaryotic genomes reveals clear species boundaries. Nat Commun 9, 5114.

JohnUrbanGenome (2022). http://biofinysics.blogspot.com/2014/05/how-does-bowtie2-assign-mapq-scores.html. Biofinysics Blog.

Kim, D., Paggi, J.M., Park, C., Bennett, C., and Salzberg, S.L. (2019). Graph-based genome alignment and genotyping with HISAT2 and HISAT-genotype. Nat Biotechnol 37, 907–915.

Kitts, P.A., Church, D.M., Thibaud-Nissen, F., Choi, J., Hem, V., Sapojnikov, V., Smith, R.G., Tatusova, T., Xiang, C., Zherikov, A., et al. (2015). Assembly: a resource for assembled genomes at NCBI. Nucleic Acids Research 44, D73–D80.

Kitts, P.A., Church, D.M., Thibaud-Nissen, F., Choi, J., Hem, V., Sapojnikov, V., Smith, R.G., Tatusova, T., Xiang, C., Zherikov, A., et al. (2016). Assembly: a resource for assembled genomes at NCBI. Nucleic Acids Res 44, D73–80.

Langmead, B., and Salzberg, S.L. (2012). Fast gapped-read alignment with Bowtie 2. Nat Methods 9, 357–359.

Langmead, B., Wilks, C., Antonescu, V., and Charles, R. (2019). Scaling read aligners to hundreds of threads on general-purpose processors. Bioinformatics 35, 421–432.

Laso-Jadart, R., Ambroise, C., Peterlongo, P., and Madoui, M.A. (2020). metaVaR: Introducing metavariant species models for reference-free metagenomic-based population genomics. PLoS One 15, e0244637.

Leggett, R.M., and MacLean, D. (2014). Reference-free SNP detection: dealing with the data deluge. BMC Genomics 15 Suppl 4, S10.

Leinonen, R., Akhtar, R., Birney, E., Bower, L., Cerdeno-Tarraga, A., Cheng, Y., Cleland, I., Faruque, N., Goodgame, N., Gibson, R., et al. (2011). The European Nucleotide Archive. Nucleic Acids Res 39, D28–31.

Leshem, A., Segal, E., and Elinav, E. (2020). The Gut Microbiome and Individual-Specific Responses to Diet. mSystems 5.

Levin, D., Raab, N., Pinto, Y., Rothschild, D., Zanir, G., Godneva, A., Mellul, N., Futorian, D., Gal, D., Leviatan, S., et al. (2021). Diversity and functional landscapes in the microbiota of animals in the wild. Science 372.

Li, H. (2018). Minimap2: pairwise alignment for nucleotide sequences. Bioinformatics 34, 3094–3100.

Li, H., and Durbin, R. (2009). Fast and accurate short read alignment with Burrows-Wheeler transform. Bioinformatics 25, 1754–1760.

Li, X., Saadat, S., Hu, H., and Li, X. (2019). BHap: a novel approach for bacterial haplotype reconstruction. Bioinformatics 35, 4624–4631.

Liu, Y., Zhang, L.Y., and Li, J. (2019). Fast detection of maximal exact matches via fixed sampling of query K-mers and Bloom filtering of index K-mers. Bioinformatics 35, 4560–4567.

Maini Rekdal, V., Bess, E.N., Bisanz, J.E., Turnbaugh, P.J., and Balskus, E.P. (2019). Discovery and inhibition of an interspecies gut bacterial pathway for Levodopa metabolism. Science 364.

Marcais, G., Delcher, A.L., Phillippy, A.M., Coston, R., Salzberg, S.L., and Zimin, A. (2018). MUMmer4: A fast and versatile genome alignment system. PLoS Comput Biol 14, e1005944.

Massana, R., and López-Escardó, D. (2022). Metagenome assembled genomes are for eukaryotes too. Cell Genomics 2.

Mitchell, C.M., Mazzoni, C., Hogstrom, L., Bryant, A., Bergerat, A., Cher, A., Pochan, S., Herman, P., Carrigan, M., Sharp, K., et al. (2020). Delivery Mode Affects Stability of Early Infant Gut Microbiota. Cell Rep Med 1, 100156.

Mukherjee, S., Seshadri, R., Varghese, N.J., Eloe-Fadrosh, E.A., Meier-Kolthoff, J.P., Goker, M., Coates, R.C., Hadjithomas, M., Pavlopoulos, G.A., Paez-Espino, D., et al. (2017). 1,003 reference genomes of bacterial and archaeal isolates expand coverage of the tree of life. Nat Biotechnol 35, 676–683.

Murray, C.S., Gao, Y., and Wu, M. (2021). Re-evaluating the evidence for a universal genetic boundary among microbial species. Nat Commun 12, 4059.

Nayfach, S., Roux, S., Seshadri, R., Udwary, D., Varghese, N., Schulz, F., Wu, D., Paez-Espino, D., Chen, I.M., Huntemann, M., et al. (2021). A genomic catalog of Earth’s microbiomes. Nat Biotechnol 39, 499–509.

Nowrotek, M., Jałowiecki, Ł., Harnisz, M., and Płaza, G.A. (2019). Culturomics and metagenomics: In understanding of environmental resistome. Frontiers of Environmental Science & Engineering 13, 40.

Olm, M.R., Crits-Christoph, A., Bouma-Gregson, K., Firek, B.A., Morowitz, M.J., and Banfield, J.F. (2021). inStrain profiles population microdiversity from metagenomic data and sensitively detects shared microbial strains. Nat Biotechnol 39, 727–736.

Olm, M.R., Crits-Christoph, A., Diamond, S., Lavy, A., Matheus Carnevali, P.B., and Banfield, J.F. (2020). Consistent Metagenome-Derived Metrics Verify and Delineate Bacterial Species Boundaries. mSystems 5.

Olson, N.D., Lund, S.P., Colman, R.E., Foster, J.T., Sahl, J.W., Schupp, J.M., Keim, P., Morrow, J.B., Salit, M.L., and Zook, J.M. (2015). Best practices for evaluating single nucleotide variant calling methods for microbial genomics. Front Genet 6, 235.

Ondov, B.D., Treangen, T.J., Melsted, P., Mallonee, A.B., Bergman, N.H., Koren, S., and Phillippy, A.M. (2016). Mash: fast genome and metagenome distance estimation using MinHash. Genome Biol 17, 132.

Ounit, R., Wanamaker, S., Close, T.J., and Lonardi, S. (2015). CLARK: fast and accurate classification of metagenomic and genomic sequences using discriminative k-mers. BMC Genomics 16, 236.

Parks, D.H., Chuvochina, M., Rinke, C., Mussig, A.J., Chaumeil, P.-A., and Hugenholtz, P. (2021). GTDB: an ongoing census of bacterial and archaeal diversity through a phylogenetically consistent, rank normalized and complete genome-based taxonomy. Nucleic Acids Research 50, D785–D794.

Parks, D.H., Rinke, C., Chuvochina, M., Chaumeil, P.A., Woodcroft, B.J., Evans, P.N., Hugenholtz, P., and Tyson, G.W. (2017). Recovery of nearly 8,000 metagenome-assembled genomes substantially expands the tree of life. Nat Microbiol 2, 1533–1542.

Peterlongo, P., Riou, C., Drezen, E., and Lemaitre, C. (2017). <em>DiscoSnp++</em>: de novo detection of small variants from raw unassembled read set(s). bioRxiv, 209965.

Phillippy, A.M., Ayanbule, K., Edwards, N.J., and Salzberg, S.L. (2009). Insignia: a DNA signature search web server for diagnostic assay development. Nucleic Acids Res 37, W229–234.

Power, R.A., Parkhill, J., and de Oliveira, T. (2017). Microbial genome-wide association studies: lessons from human GWAS. Nat Rev Genet 18, 41–50.

Pulido-Tamayo, S., Sanchez-Rodriguez, A., Swings, T., Van den Bergh, B., Dubey, A., Steenackers, H., Michiels, J., Fostier, J., and Marchal, K. (2015). Frequency-based haplotype reconstruction from deep sequencing data of bacterial populations. Nucleic Acids Res 43, e105.

R Core Development Team, T. (2022). The R Project for Statistical Computing: https://www.r-project.org.

Rodriguez, R.L., Jain, C., Conrad, R.E., Aluru, S., and Konstantinidis, K.T. (2021). Reply to: “Re-evaluating the evidence for a universal genetic boundary among microbial species”. Nat Commun 12, 4060.

Saak, C.C., Dinh, C.B., and Dutton, R.J. (2020). Experimental approaches to tracking mobile genetic elements in microbial communities. FEMS Microbiol Rev 44, 606–630.

Sarhan, M.S., Hamza, M.A., Youssef, H.H., Patz, S., Becker, M., ElSawey, H., Nemr, R., Daanaa, H.A., Mourad, E.F., Morsi, A.T., et al. (2019). Culturomics of the plant prokaryotic microbiome and the dawn of plant-based culture media - A review. J Adv Res 19, 15–27.

Schloissnig, S., Arumugam, M., Sunagawa, S., Mitreva, M., Tap, J., Zhu, A., Waller, A., Mende, D.R., Kultima, J.R., Martin, J., et al. (2013). Genomic variation landscape of the human gut microbiome. Nature 493, 45–50.

Shah, R.N., and Ruthenburg, A.J. (2021). Sequence deeper without sequencing more: Bayesian resolution of ambiguously mapped reads. PLoS Comput Biol 17, e1008926.

Shajii, A., Yorukoglu, D., William Yu, Y., and Berger, B. (2016). Fast genotyping of known SNPs through approximate k-mer matching. Bioinformatics 32, i538–i544.

Shi, Z.J., Dimitrov, B., Zhao, C., Nayfach, S., and Pollard, K.S. (2022). Fast and accurate metagenotyping of the human gut microbiome with GT-Pro. Nat Biotechnol 40, 507–516.

Shoemaker, W.R., Chen, D., and Garud, N.R. (2022). Comparative Population Genetics in the Human Gut Microbiome. Genome biology and evolution 14.

Smillie, C.S., Sauk, J., Gevers, D., Friedman, J., Sung, J., Youngster, I., Hohmann, E.L., Staley, C., Khoruts, A., Sadowsky, M.J., et al. (2018). Strain Tracking Reveals the Determinants of Bacterial Engraftment in the Human Gut Following Fecal Microbiota Transplantation. Cell Host Microbe 23, 229–240 e225.

Smits, S.A., Leach, J., Sonnenburg, E.D., Gonzalez, C.G., Lichtman, J.S., Reid, G., Knight, R., Manjurano, A., Changalucha, J., Elias, J.E., et al. (2017). Seasonal cycling in the gut microbiome of the Hadza hunter-gatherers of Tanzania. Science 357, 802–806.

Sood, U., Kumar, R., and Hira, P. (2021). Expanding Culturomics from Gut to Extreme Environmental Settings. mSystems, e0084821.

Truong, D.T., Franzosa, E.A., Tickle, T.L., Scholz, M., Weingart, G., Pasolli, E., Tett, A., Huttenhower, C., and Segata, N. (2015). MetaPhlAn2 for enhanced metagenomic taxonomic profiling. Nat Methods 12, 902–903.

Vainberg-Slutskin, I., Kowalsman, N., Silberberg, Y., Cohen, T., Gold, J., Kario, E., Weiner, I., Gahali-Sass, I., Kredo-Russo, S., Zak, N.B., et al. (2022). Exodus: sequencing-based pipeline for quantification of pooled variants. Bioinformatics.

Van Rossum, T., Costea, P.I., Paoli, L., Alves, R., Thielemann, R., Sunagawa, S., and Bork, P. (2021). metaSNV v2: detection of SNVs and subspecies in prokaryotic metagenomes. Bioinformatics.

Van Rossum, T., Ferretti, P., Maistrenko, O.M., and Bork, P. (2020). Diversity within species: interpreting strains in microbiomes. Nat Rev Microbiol 18, 491–506.

Xie, H., Yang, C., Sun, Y., Igarashi, Y., Jin, T., and Luo, F. (2020). PacBio Long Reads Improve Metagenomic Assemblies, Gene Catalogs, and Genome Binning. Front Genet 11, 516269.

Yahara, K., Suzuki, M., Hirabayashi, A., Suda, W., Hattori, M., Suzuki, Y., and Okazaki, Y. (2021). Long-read metagenomics using PromethION uncovers oral bacteriophages and their interaction with host bacteria. Nat Commun 12, 27.

Zeevi, D., Korem, T., Godneva, A., Bar, N., Kurilshikov, A., Lotan-Pompan, M., Weinberger, A., Fu, J., Wijmenga, C., Zhernakova, A., et al. (2019). Structural variation in the gut microbiome associates with host health. Nature 568, 43–48.

Zeng, Q., Liao, C., Terhune, J., and Wang, L. (2019). Impacts of florfenicol on the microbiota landscape and resistome as revealed by metagenomic analysis. Microbiome 7, 155.

Zhao, C., Dimitrov, B., Goldman, M., Nayfach, S., and Pollard, K.S. (2022). MIDAS2: Metagenomic Intra-species Diversity Analysis System. bioRxiv, 2022.2006.2016.496510.

Zheng, Y., Ay, F., and Keles, S. (2019). Generative modeling of multi-mapping reads with mHi-C advances analysis of Hi-C studies. Elife 8.

